# Uncoupling mycomembrane biogenesis from mycolic acid synthesis reveals a distinct role for mycoloyltransferases in mycobacterial cell division

**DOI:** 10.64898/2026.07.20.738443

**Authors:** Célia de Sousa-d’Auria, Florence Constantinesco-Beker, Magali Prigent, Fatima Taiki, Claire Boulogne, Louis-David Leclercq, Lena Rosier, Mickaël Bourge, Vlad Costache, Mohamed Chami, Cécile Labarre, Christine Houssin, Anne Marie Wehenkel, Amélie Leforestier, Yann Guérardel, Nicolas Bayan

## Abstract

Bacteria of the order Mycobacteriales, including the genera *Mycobacterium* and *Corynebacterium*, possess a unique outer membrane, termed the mycomembrane, which is structurally and chemically distinct from the lipopolysaccharide-containing outer membrane of Gram-negative bacteria. A defining feature of the mycomembrane is its enrichment in mycolic acids, long-chain α-branched, β-hydroxylated fatty acids that occur as trehalose monomycolate (TMM), trehalose dimycolate (TDM), or are esterified to arabinogalactan, an unusual polymer that is itself covalently linked to peptidoglycan (PG). Mycoloyltransferases are Mycobacteriales*-*specific enzymes described to catalyze the transfer of mycolic acids from trehalose monomycolate (TMM) to various cell envelope acceptors, including trehalose and arabinogalactan. In several species, including *Mycobacterium tuberculosis*, these proteins are essential for viability; however, their occurrence as multiple paralogs with partially redundant functions has hindered the precise assignment of their cellular roles.

Previously, we showed that *Corynebacterium glutamicum* remains viable in the absence of mycolic acids, and thus without a mycomembrane, following deletion of *pks*, the gene required for mycolic acid biosynthesis. Building on this finding, we systematically deleted all genes encoding mycoloyltransferases in *C. glutamicum* to further disclose their collective function in the cell. The resulting Δ*myts* mutant lacked arabinogalactan-bound mycolates and TDM, and continued to synthesize TMM. While this major outer membrane glycolipid is highly abundant, the Δ*myts* strain failed to assemble a mycomembrane and displayed pronounced cell aggregation. Deletion of mycoloyltransferases also caused very severe defects in cell division and morphogenesis that are not observed in a Δ*pks* strain unable to synthesize mycolic acids. Together, these results demonstrate that mycoloyltransferases are essential for mycomembrane assembly but dispensable for TMM biosynthesis, and reveal an important role for these enzymes in cell division that is independent of their canonical mycolic acid transfer activity.

**SIGNIFICANCE:** How the mycomembrane is assembled and anchored to the cell wall remains a central question in Mycobacteriales, where this outer membrane is necessary for envelope integrity and intrinsic antibiotic resistance. By genetically separating mycolic acid synthesis from their incorporation into the outer membrane, we identify arabinogalactan-linked mycolates as the critical determinant for initiating membrane assembly. Importantly, we also uncover a role for mycoloyltransferases beyond their canonical function in lipid metabolism, revealing a functional link with cell division. These findings point to a critical role of Mycoloyltransferases in the coordination between outer membrane biogenesis and bacterial cytokinesis.

## INTRODUCTION

Members of the suborder Mycobacteriales, which includes *Mycobacterium tuberculosis* and *Corynebacterium glutamicum*, possess a highly specialized cell envelope that is essential for their physiology and environmental resilience. Although taxonomically classified as Gram-positive, these bacteria display a didermic envelope organization featuring a distinctive outer membrane known as the mycomembrane [1]. This structure is largely composed of lipids containing mycolic acids (long-chain, α-branched, β-hydroxylated fatty acids), that confer effective protection against hydrolytic damage. The synthesis of these unique fatty acids begins in the cytoplasm where individual acyl chains are condensed by a specific cytoplasmic polyketide synthase (Pks in *C. glutamicum* or Pks13 in *M. tuberculosis*)[2], [3]. Based on *in vitro* studies, the cytoplasmic polyketide synthase Pks13 has also been proposed to catalyze the subsequent transfer of mycolate chains onto trehalose. The newly born trehalose monomycolate (TMM) is then flipped from the inner to the outer leaflet of the plasma membrane by an RND transporter of the MmpL family [4]. Finally, the mycolate chain of the exported TMM is redistributed to various mycomembrane building blocks (including some proteins) by a family of enzymes called mycoloyltransferases (Myts) [5], [6], [7], [8], [9]. These enzymes are homologous α/β-hydrolase fold proteins highly conserved in Mycobacteriales and that include the previously characterized antigen 85 complex of *M. tuberculosis* [10], [11], [12]. They are exported to the periplasm and are found both associated to the cell envelope and secreted in the culture medium [11], [13]. They mediate mycoloyl transfer reactions, transferring mycolic acids from TMM to various acceptors. The main reactions include mycoloylation of TMM to form trehalose dimycolate (TDM) and the mycoloylation of the multiple terminal arabinofuranosyl motif of arabinogalactan (AG) polymer (AG-bound mycolates), which links the mycomembrane to the underlying peptidoglycan meshwork. All Mycobacteriales species contains several paralogs (from 3 to 11) sharing conserved catalytic motifs but with similarity scores ranging from 30 to 80%, suggesting that although they have similar catalytic activity, they may have distinct functions in the cell.

In *Mycobacterium* species, three catalytically active mycoloyltransferases, Ag85A, Ag85B, and Ag85C, have been extensively characterized. Genetic inactivation of *ag85C* in *M. tuberculosis* results in a marked reduction (∼40%) of cell-wall-bound mycolates and increased envelope permeability, underscoring the importance of individual Myts for envelope integrity [6]. In addition to its canonical mycoloyltransferase activity, Ag85A was shown, upon overexpression, to possess acyl-CoA:diacylglycerol acyltransferase (DGAT) activity [14], suggesting a potential role in triacylglycerol accumulation and bacterial dormancy and highlighting the functional versatility of this protein family. Although double *ag85* mutants have been generated [15], a viable mutant lacking all three active Ag85 mycolyltransferases has not been reported, suggesting that their combined activity may be essential for viability and making them attractive targets for antimicrobial development. A fourth paralog, Ag85D, is present in several *Mycobacterium* species but lacks key catalytic residues, and its physiological role remains unknown [16].

In *Corynebacterium glutamicum*, a genetically tractable nonpathogenic model organism for Mycobacteriales, six *myt* genes (*mytA–F*) have been identified [7]. Early studies demonstrated that inactivation of *mytA* caused a substantial reduction in cell-wall-bound mycolates and TDM, accompanied by accumulation of TMM, establishing MytA as a major mycoloyltransferase required for both arabinogalactan- and trehalose-linked mycolate formation [8]. Subsequent work [7] identified additional Myt paralogs and classified them into three functional groups (MytA/B, MytD/E/F, and MytC). While MytA/B and MytD/E/F exhibited partial redundancy in TDM synthesis and arabinogalactan mycoloylation, MytC was shown to catalyze O-mycoloylation of specific exported proteins [17], [18] rather than participating directly in mycomembrane assembly. Disruption of multiple *myt* genes, particularly *mytA* and *mytB*, severely reduced cell-wall-bound corynomycolates, abolished the outer electron-dense layer of the envelope, and resulted in temperature-sensitive growth and pronounced cell-separation defects [9], [19].

Collectively, these studies indicate that the presence of multiple mycoloyltransferase paralogs is essential to coordinate mycolate transfer to diverse acceptors, yet the extensive functional redundancy among Myts has hindered a system-level understanding of how mycoloyltransferase activity integrates mycolic acid synthesis with mycomembrane assembly and cell growth. Here, to address this question and reveal mycoloyltransferase functions beyond individual substrate specificity, we choose a global genetic approach and decided to construct a *C. glutamicum* strain lacking all six known mycoloyltransferases (Δ*myts*). Our results show that this mutant was unable to produce arabinogalactan-bound mycolates or trehalose dimycolate, but still retain the ability to synthesize and export TMM, demonstrating that TMM biosynthesis occurs completely independently of Myt activity. Despite continued TMM production, the mutant failed to assemble a mycomembrane, indicating that TMM production alone is insufficient for mycomembrane biogenesis and highlighting the essential role of mycoloyltransferases in coupling mycolic acid incorporation to envelope assembly. Loss of Myt activity also resulted in severe defects in cell morphology and division. These defects persisted in a Δ*pks* background, despite the absence of morphological defects in the Δ*pks* mutant itself, indicating that they cannot be attributed to the loss of the mycomembrane. These findings reveal an unanticipated role for mycoloyltransferases in coordinating cell envelope biogenesis with cell division and position Myts as central regulators of cell growth beyond their canonical enzymatic function.

## RESULTS

### The Δ*myts* mutant is viable but displays significant growth defect

In *Corynebacterium glutamicum*, genes encoding mycoloyltransferases are distributed across the genome (Fig. S1). Notably, *mytA* and *mytB* are located adjacent to each other, separated only by a small ORF, within an ∼21-kb region that is highly conserved among Mycobacteriales [20]. This locus also contains genes involved in cell envelope biogenesis, including *pccB-pks-fadD2* for mycolic acid synthesis and *aftB* or *glfT* for arabinogalactan biosynthesis (Fig. S1). To investigate the role of mycoloyltransferases, we sequentially deleted each of the *myt* genes in the following order: *mytF*, *mytE*, *mytD*, *mytC*, *mytA*, and *mytB*, using the method described by Schäfer et al.[21], which ensures complete gene deletion without leaving a resistance cassette. For this end, a pkdel*myt* deletion vector was constructed for each gene, and a series of mutants was generated to obtain a sextuple mutant, designated Δ*myts* (Table 1).

**Table 1:**
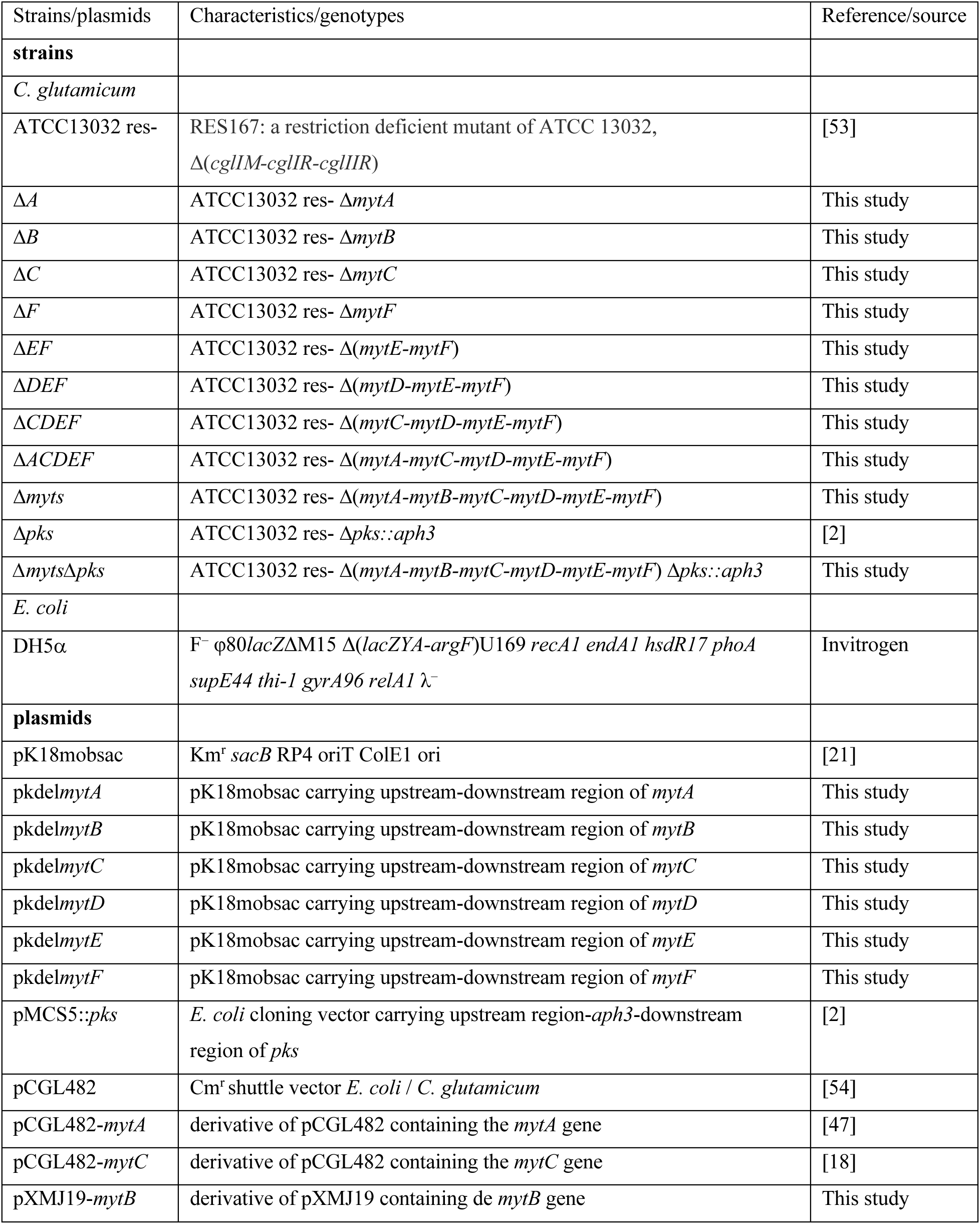
Strains and plasmids used in this study.

| Strains/plasmids | Characteristics/genotypes | Reference/source |
|---|---|---|
| **strains** |  |  |
| *C. glutamicum* |  |  |
| ATCC13032 res- | RES167: a restriction deficient mutant of ATCC 13032, Δ(*cglIM-cglIR-cglIIR*) | [53] |
| Δ*A* | ATCC13032 res- Δ*mytA* | This study |
| Δ*B* | ATCC13032 res- Δ*mytB* | This study |
| Δ*C* | ATCC13032 res- Δ*mytC* | This study |
| Δ*F* | ATCC13032 res- Δ*mytF* | This study |
| Δ*EF* | ATCC13032 res- Δ(*mytE-mytF*) | This study |
| Δ*DEF* | ATCC13032 res- Δ(*mytD-mytE-mytF*) | This study |
| Δ*CDEF* | ATCC13032 res- Δ(*mytC-mytD-mytE-mytF*) | This study |
| Δ*ACDEF* | ATCC13032 res- Δ(*mytA-mytC-mytD-mytE-mytF*) | This study |
| Δ*myts* | ATCC13032 res- Δ(*mytA-mytB-mytC-mytD-mytE-mytF*) | This study |
| Δ*pks* | ATCC13032 res- Δ*pks::aph3* | [2] |
| Δ*myts*Δ*pks* | ATCC13032 res- Δ(*mytA-mytB-mytC-mytD-mytE-mytF*) Δ*pks::aph3* | This study |
| *E. coli* |  |  |
| DH5α | F<sup>–</sup> φ80*lacZ*ΔM15 Δ(*lacZYA-argF*)U169 *recA1 endA1 hsdR17 phoA supE44 thi-1 gyrA96 relA1* λ<sup>–</sup> | Invitrogen |
| **plasmids** |  |  |
| pK18mobsac | Km<sup>r</sup> *sacB* RP4 oriT ColE1 ori | [21] |
| pkdel*mytA* | pK18mobsac carrying upstream-downstream region of *mytA* | This study |
| pkdel*mytB* | pK18mobsac carrying upstream-downstream region of *mytB* | This study |
| pkdel*mytC* | pK18mobsac carrying upstream-downstream region of *mytC* | This study |
| pkdel*mytD* | pK18mobsac carrying upstream-downstream region of *mytD* | This study |
| pkdel*mytE* | pK18mobsac carrying upstream-downstream region of *mytE* | This study |
| pkdel*mytF* | pK18mobsac carrying upstream-downstream region of *mytF* | This study |
| pMCS5::*pks* | *E. coli* cloning vector carrying upstream region-*aph3*-downstream region of *pks* | [2] |
| pCGL482 | Cm<sup>r</sup> shuttle vector *E. coli* / *C. glutamicum* | [54] |
| pCGL482-*mytA* | derivative of pCGL482 containing the *mytA* gene | [47] |
| pCGL482-*mytC* | derivative of pCGL482 containing the *mytC* gene | [18] |
| pXMJ19-*mytB* | derivative of pXMJ19 containing de *mytB* gene | This study |

To assess the functional consequences of losing all mycoloyltransferase activity, we compared Δ*myts* to the wild-type (WT) strain and to the Δ*pks* mutant, which is deficient in mycolic acid synthesis. Phenotypically, Δ*myts* formed matte, rough colonies resembling Δ*pks*, in contrast to the smooth, glossy appearance of the wild type (WT) (Fig. 1A). In liquid culture, Δ*myts* grew slowly and formed large aggregates at 30°C, similar to Δ*pks* (Fig. 1B). To accurately monitor growth, cultures were shaken in baffled flasks to limit aggregation and allow reliable optical density measurements at 30°C. Results from three independent experiments (Fig. 1C) showed that Δ*myts* displayed a growth delay comparable to Δ*pks*, but with greater variability between replicates. The average doubling times were 57 ± 2 min for WT, 123 ± 4 min for Δ*pks*, and 138 ± 27 min for Δ*myts*.

**Figure 1:**
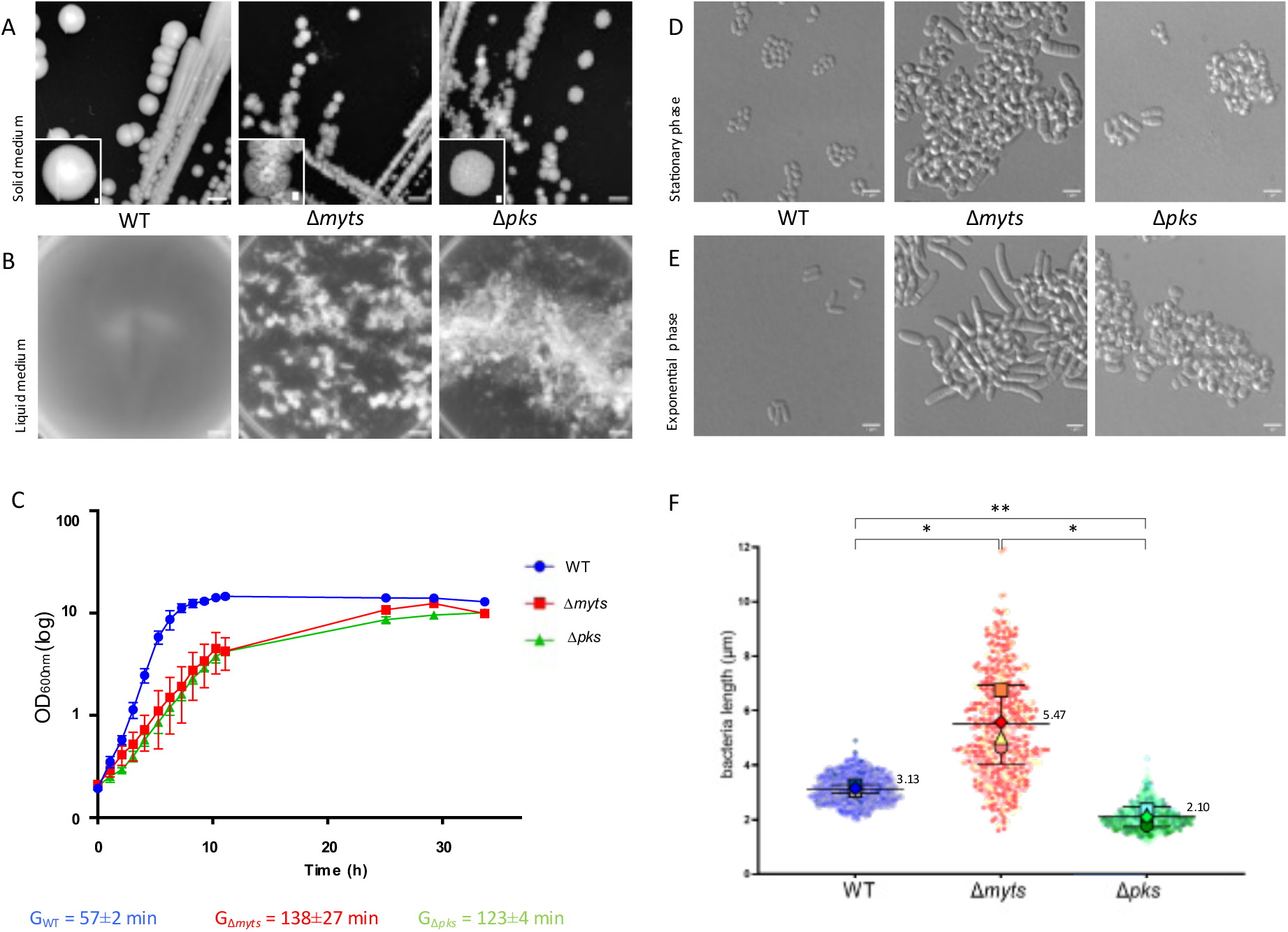
Morphology and growth characteristics of WT, Δ*myts*, and Δ*pks* strains. **(A)** Colony morphology on solid BHI medium (Arbitrary scale). **(B)** Appearance of cultures grown in liquid BHI medium for 24 h at 30°C (Arbitrary scale). **(C)** Growth curves of cells grown in BHI medium at 30°C in baffled shake flasks. Curves represent the mean of three independent replicates; individual data points are shown (WT, blue circles; Δ*myts*, red squares; Δ*pks*, green triangles). Generation time (G) for each strain is indicated just below the graph. **(D,E)** Representative differential interference contrast (DIC) micrographs of cells grown in BHI medium at 25°C during stationary phase **(D)** and exponential phase **(E)**. Scale bars, 3 µm. **(F)** SuperPlots comparing cell length distributions of WT, Δ*myts*, and Δ*pks* strains during exponential growth at 25 °C. Cell lengths were measured from four independent biological replicates per strain. Small circles represent individual cells and are color-coded by replicate (WT, shades of blue; Δ*myts*, shades of red; Δ*pks*, shades of green). Large symbols indicate the mean of each biological replicate (square, replicate 1; circle, replicate 2; diamond, replicate 3; triangle, replicate 4), shown in the corresponding strain color and used to present summary descriptive statistics: the overall mean is indicated by a large bar with error bars (95% confidence interval). Cell length differences between populations were significant within each biological replicate **(Fig. S2)**. However, while the WT showed consistent cell lengths across independent experiments, the Δ*myts* and Δ*pks* mutants displayed greater variability between replicates **(Table S3)**. To account for this variability, statistical analyses of the superplots were performed using Welch’s ANOVA followed by Dunnett’s T3 post-hoc test. Significant comparisons are marked by asterisks. *, P < 0.05; **, P < 0.01 **(Table S3)**.

Microscopic examination revealed that Δ*myts* cells were highly elongated and morphologically irregular during both stationary and exponential phases (Fig. 1D and 1E), whereas Δ*pks* cells appeared wider and more coccoid than the wild type (WT). Quantitative analysis of exponential-phase cultures from four independent experiments (Fig. 1F and Fig. S2, Table S2) confirmed significant heterogeneity in Δ*myts* cell length, which ranged from 1.61 µm to 11.93 µm (mean = 5.47 µm). This distribution starkly contrasts with the lengths measured for both the WT (1.96–4.90 µm, mean = 3.13 µm) and the Δ*pks* populations (1.11–4.24 µm, mean = 2.10 µm). Significant differences in cell length distributions were observed among all three strains in each experiment (Fig. S2; Table S2 and S3). However, Δ*myts* and Δ*pks* exhibited greater inter-experimental variability than the WT.

Interestingly, examination of the intermediate mutants generated during the sequential deletion of the *myt* genes revealed that changes in cell aggregation and morphology became progressively apparent upon deletion of *mytC (ΔCDEF)*, *mytA (ΔACDEF)*, and *mytB (Δmyts)*, the last three genes deleted during the construction of the Δ*myts* mutant (Fig. S3). This observation suggested that MytA, MytB, and MytC, which are also the most expressed Myts in the cell, make major contributions to maintaining normal cell morphology and limiting aggregation. We therefore further characterized the corresponding single-deletion mutants (Δ*mytA*, Δ*mytB*, and Δ*mytC*), all of which displayed morphological defects of varying severity (Fig. S4). Notably, Δ*mytA* cells appear markedly elongated, whereas Δ*mytB* cells exhibit a more coccoid morphology. Δ*mytC* cells exhibited a morphology largely comparable to that of the wild type, with the exception of occasional isolated, unusually elongated cells.

### Δ*myts* synthesizes TMM but fails to assemble a mycomembrane

Because mycoloyltransferases are involved in the formation of both AG-bound mycolates and trehalose mycolates, we analyzed the lipid composition of the Δ*myts* mutant. First, total lipid fractions were extracted from stationary-phase cultures and analyzed by TLC, revealing differential accumulation of glycolipid species among the strains (Fig. 2A). The identities of the major mycolate-containing glycolipids, TMM (Rf = 0,46) and TDM (Rf = 0,92), were subsequently confirmed by purification and structural analyses using mass spectrometry and NMR (Fig. S5-S6, Table S4). As expected, the Δ*pks* strain lacked all mycolates derived component (Fig. 2A and 2B). In contrast, the Δ*myts* failed to produce trehalose dimycolate (TDM) and AG-bound mycolates, but accumulated trehalose monomycolate (TMM). In addition, Δ*myts* cells seems to contain lower levels of phosphatidylinositol mannosides (PIMs) than WT cells. Analysis of the culture supernatant by high-speed centrifugation revealed lipid-like material in the pellet, predominantly composed of TMM (Fig. 2C) but also small amounts of phosphatidylinositol (PI) and PIMs. Further purification of this material on a sucrose cushion followed by cryo–electron microscopy confirmed that it is composed of extracellular vesicles of various sizes but also of undefined structures ressembling large sheets or open vesicles. In contrast to vesicles produced by the Δ*aftB* mutant, which releases mycomembrane-derived vesicles containing both TDM and TMM [22], these vesicles do not contain TDM but almost exclusively TMM (Fig. 2C) and small amounts of PI and PIMs suggesting they may originate, at least in part, from the inner membrane.

**Figure 2:**
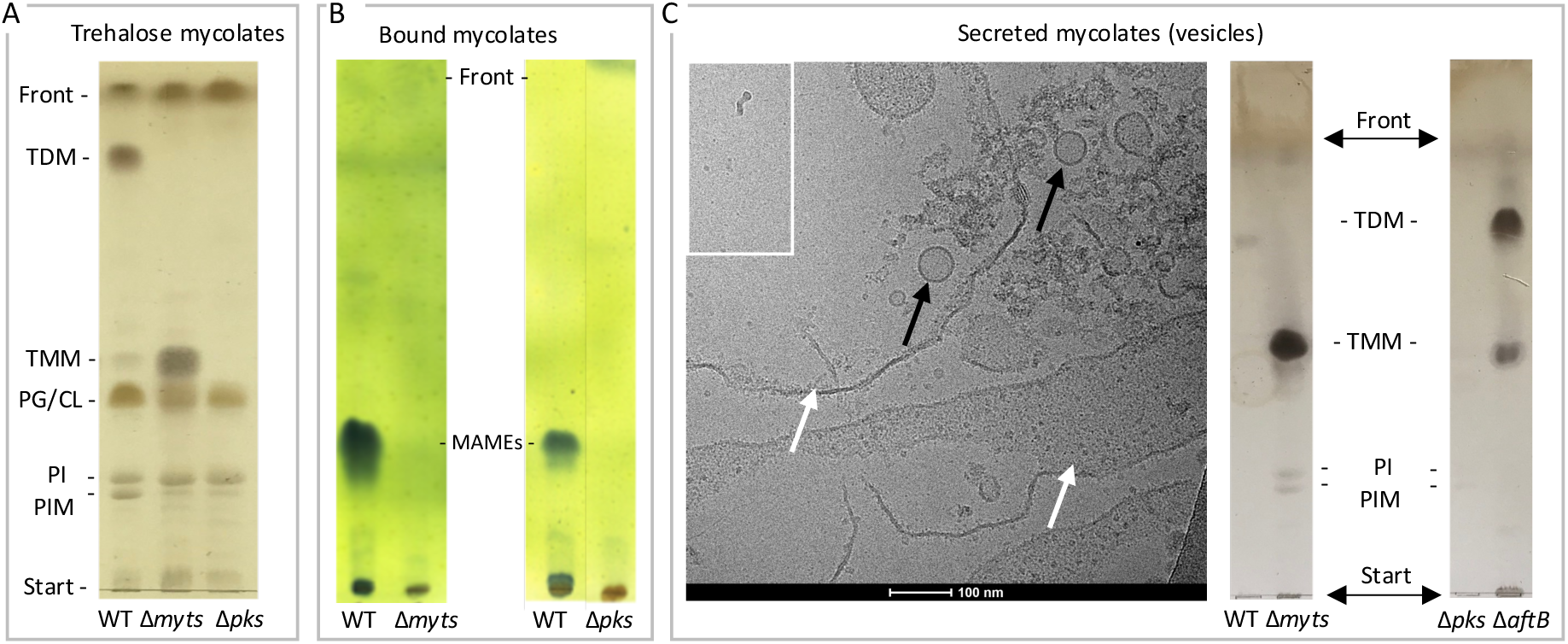
Lipid analysis of the Δ*myts* mutant. **(A, B**) Thin-layer chromatography (TLC) analysis of lipid extracts from whole cells of WT, Δ*myts*, and Δ*pks* strains. Extracts containing **(A)** trehalose mycolates or **(B)** bound mycolates (esterified to arabinogalactan) were prepared and resolved using appropriate solvent systems as described in Materials and Methods. The nature of trehalose mycolates TMM and TDM were confirmed by MS and NMR (Fig. S5 and S6). **(C)** Analysis of material released into the culture supernatant by the Δ*myts* mutant. Vesicles (black arrows) or undefined material, large sheets or open vesicles, (white arrows) are observed. Ultracentrifugation of culture supernatant from Δ*myts* revealed a clearly visible pellet (shown in the **inset of C**). This material was further analyzed by cryo-electron microscopy and TLC. Δ*aftB* secreted vesicles previously described by Bou Raad *et al.* [22] were also prepared and included for comparison in this figure. The migration positions of reference lipids are indicated: TDM, trehalose dimycolate (Rf= 0,92); TMM, trehalose monomycolate (Rf = 0,46); PG/CL, phosphatidylglycerol/cardiolipin (Rf = 0,39); PI, phosphatidylinositol (Rf = 023); PIM, phosphatidylinositol mannoside (Rf = 0,2); MAMEs, mycolic acids methyl esters.

To assess whether the Δ*myts* strain retained an outer mycomembrane despite the absence of AG-bound mycolates, we first used a mycomembrane-specific fluorescent probe: a rhodamine-labeled phospholipid analog, rDHPE (2-dihexadecanoyl-sn-glycero-3-phosphoethanolamine) [23]. Cells were also stained with BODIPY FL–vancomycin (Van-FL) to visualize nascent peptidoglycan [24]. Representative fluorescence images are shown in Fig. S7. In WT cells, Van-FL staining was strongest at the midcell septum, and at the poles of newly separated cells. rDHPE labeled the entire WT cell surface uniformly, occasionally showing stronger fluorescence at the septum. Some Van-FL-positive septa were not stained by rDHPE, consistent with previous reports indicating that mycolic acid layers form after the peptidoglycan layer [25], [26]. In Δ*pks* cells, two morphotypes were observed: elongated cells with one or two Van-FL-positive septa of similar intensity to the lateral envelope, and shorter cells displaying intense polar or patchy surface Van-FL labeling (white arrows of Fig. S7). Brightly stained bud-like structures were also detected (blue arrows of Fig. S7). As expected, in both morphotypes, rDHPE labeling is not detected. In the Δ*myts* mutant, multiple Van-FL-positive septa were visible within individual cells. Septal labeling intensity varied, with localized brighter regions along the same septum (yellow arrows on Δ*myts* of Fig. S7). As observed for Δ*pks*, the Δ*myts* strain showed no detectable rDHPE labeling, indicating a severe disruption or absence of the mycomembrane.

To confirm this observation at the ultrastructural level, we used cryo-electron microscopy of vitreous sections (CEMOVIS), a reference method that initially demonstrated the lack of a mycomembrane in the *C. glutamicum* Δ*pks* mutant [27]. This technique involves high-pressure freezing followed by cryosectioning of vitrified, fully hydrated samples for observation by cryo-electron microscopy. CEMOVIS images and corresponding density profiles (Fig. 3) clearly revealed the absence of a mycomembrane in both Δ*pks* and Δ*myts* mutants. The measured envelope thicknesses of WT, Δ*myts* and Δ*pks* cells is reported in Fig. 3G and the procedures of measurement in Fig. S8.

**Figure 3:**
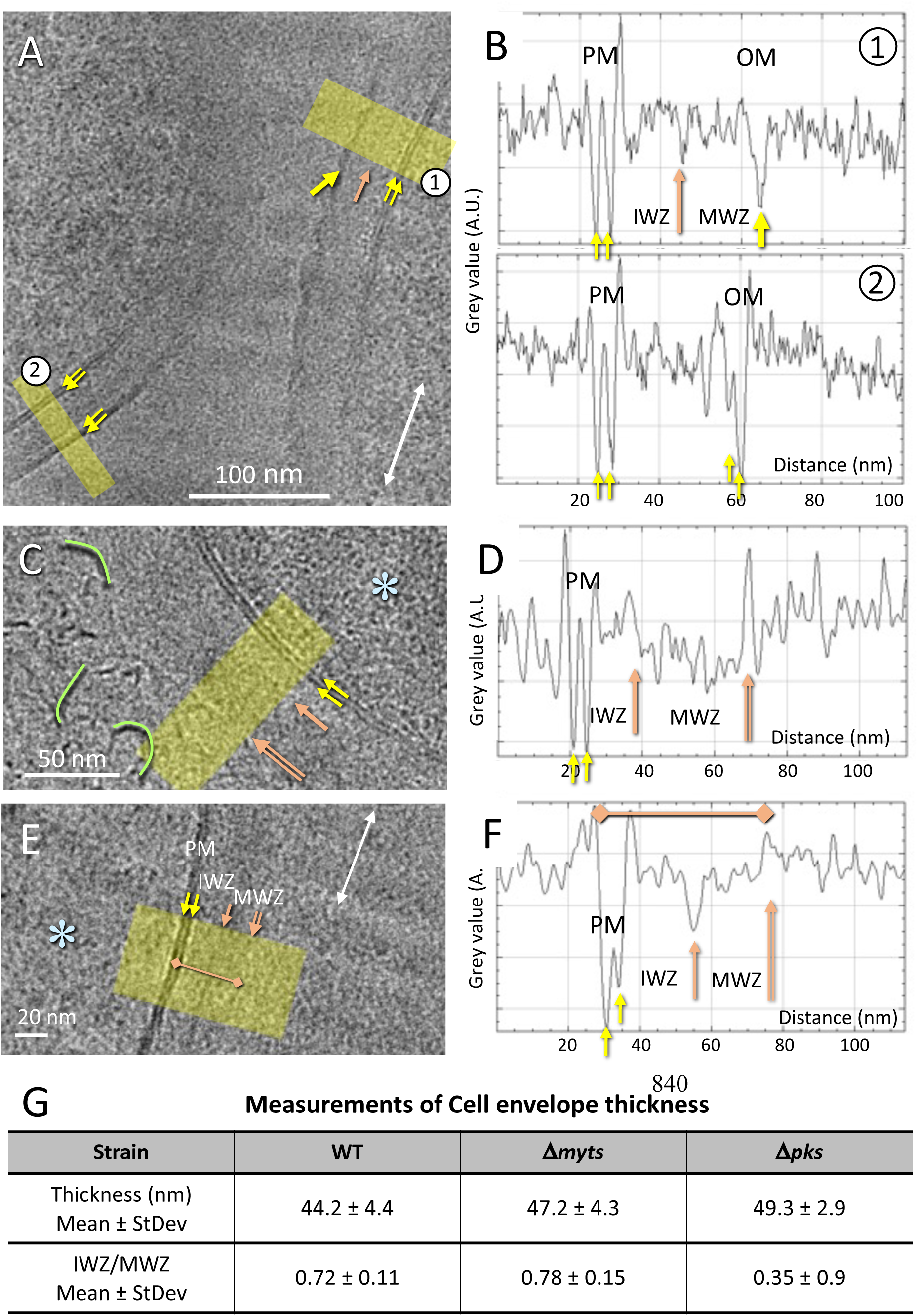
CEMOVIS observations of WT, Δ*myts* and Δ*pks* strains. **(A, B1, B2)** WT cells present a typical architecture envelope with their CW between two lipid bilayers. Our data confirm previous observations of Zuber *et al.*[27], in particular the existence of an outer membrane (OM) with its bilayer leaflet. It is thinner than that of the plasma membrane (PM) and appears destabilized in many places (line profile (1) and thick yellow arrow), a phenomenon also observed by Zuber *et al*. [27]. The two leaflets of bilayers are indicated by two parallel yellow arrows. **(C)** Δ*pks* cell envelope detail, with the plasma membrane (PM) double leaflet (yellow arrows), the inner wall zone (IWZ) and the median wall zone (MWZ). Some filaments filling the external medium are underlined in acid green. **(D)** line profile from (**C**, yellow overlay) with corresponding peaks. **(E)** Detail of Δ*myts* cell envelope, with the plasma membrane (PM) double leaflet (yellow arrows), the inner wall zone (IWZ) and the median wall zone (MWZ). **(F)** Line profile from (E, yellow overlay) with corresponding peaks. In **(A, C, E),** the white double arrow underlines knife marks on the section’s surface, highlighting the cutting direction. The envelope dimensions measured in areas where the cell envelope is parallel or close to the cutting direction to avoid sectioning-induced compression effects are shown in **G** (see details in Figure S6). **(A, C, E)** CEMOVIS images. **(B1, B2, D, F)** density line profiles. Glacial blue asterisks indicate cytoplasm. **(G)** Measurements of total cell envelope thickness and IWZ/MWZ thickness ratio.

### Δ*myts* cells exhibit dramatic morphological defects

In order to more carefully analyze the morphology defect of our Δ*myts* mutant, detected by brightfield microscopy, we decided to complement our studies with SEM, CEMOVIS and FIB-SEM observations that are complementary techniques to obtain higher-resolution information on both surface architecture and internal organization both at individual and population levels. SEM imaging of the three strains in exponential and stationary phases (Fig. S9) revealed striking differences in morphology and population structure. While WT cells appear as relatively uniform rods forming small clusters, the Δ*myts* population is highly heterogeneous, containing short cells alongside markedly elongated and distorted forms that assemble into irregular aggregates. In contrast, the Δ*pks* mutant, though stockier and much more homogenous, also forms large aggregates. Clearly, the Δ*myts* strain reveals some phenotypic characteristics pointing toward a dramatic cell division defect that is not observed in the Δ*pks* strain and therefore not related to the absence of the mycomembrane which is absent in both strains. In-depth analyses of Δ*myts* cells compared to WT and Δ*pks* cells using CEMOVIS and FIB-SEM confirm and further reveal striking morphological differences. Representative images of the Δ*myts* cell division phenotype are shown in Figure 4 (and enlarged in Fig. S10) with most abnormal morphologies visualized in parallel by SEM, FIB-SEM and CEMOVIS. A similar synthesis is shown for the WT and Δ*pks* strains for comparison (Fig. S11 and Fig. S12). As it is clearly visible on FIB-SEM images (Fig. 4, Fig. S13-14 and video 3), Δ*myts* cells are notably elongated and clearly shows the presence of multiple septa within individual cells, frequently positioned abnormally: curved, misaligned, closely spaced, or even intersecting (Fig. 4 and Fig. S7B, Fig.S13). Some of them are fully formed and others develop across segregating nucleoids (Fig. S14 and Video 4), a division phenotype previously described in *Mycobacterium*, and *Corynebacterium* [28]. Many septa appear curved or dome-shaped, consistent with continued polar growth despite failed daughter-cell separation, resulting in the accumulation of multiple nascent daughter compartments within individual Δ*myts* cells (blue asterisks, Fig. 4). Rather than undergoing controlled V-snapping, these compartments frequently appear to rupture, leading to the release of the contents of the enclosed cells (orange asterisks), often in association with vesicle formation (yellow arrows). These damaged regions are readily identifiable in FIB-SEM as seam-like openings oriented perpendicular to the septa (Fig. S14 and video 4), a feature also evident in SEM (Fig. 4). Vesicle production is frequently observed in both SEM and FIB-SEM, consistent with the formation of empty or “ghost” cell ends seen in SEM and CEMOVIS. More FIB-SEM images (Fig. S13) are shown in supplementary material to have better representative view of the global differences between the three strains. A synthesis of the main characteristics observed by electron microscopy (SEM, FIB-SEM, CEMOVIS) is recapitulated in Table S5.

**Figure 4:**
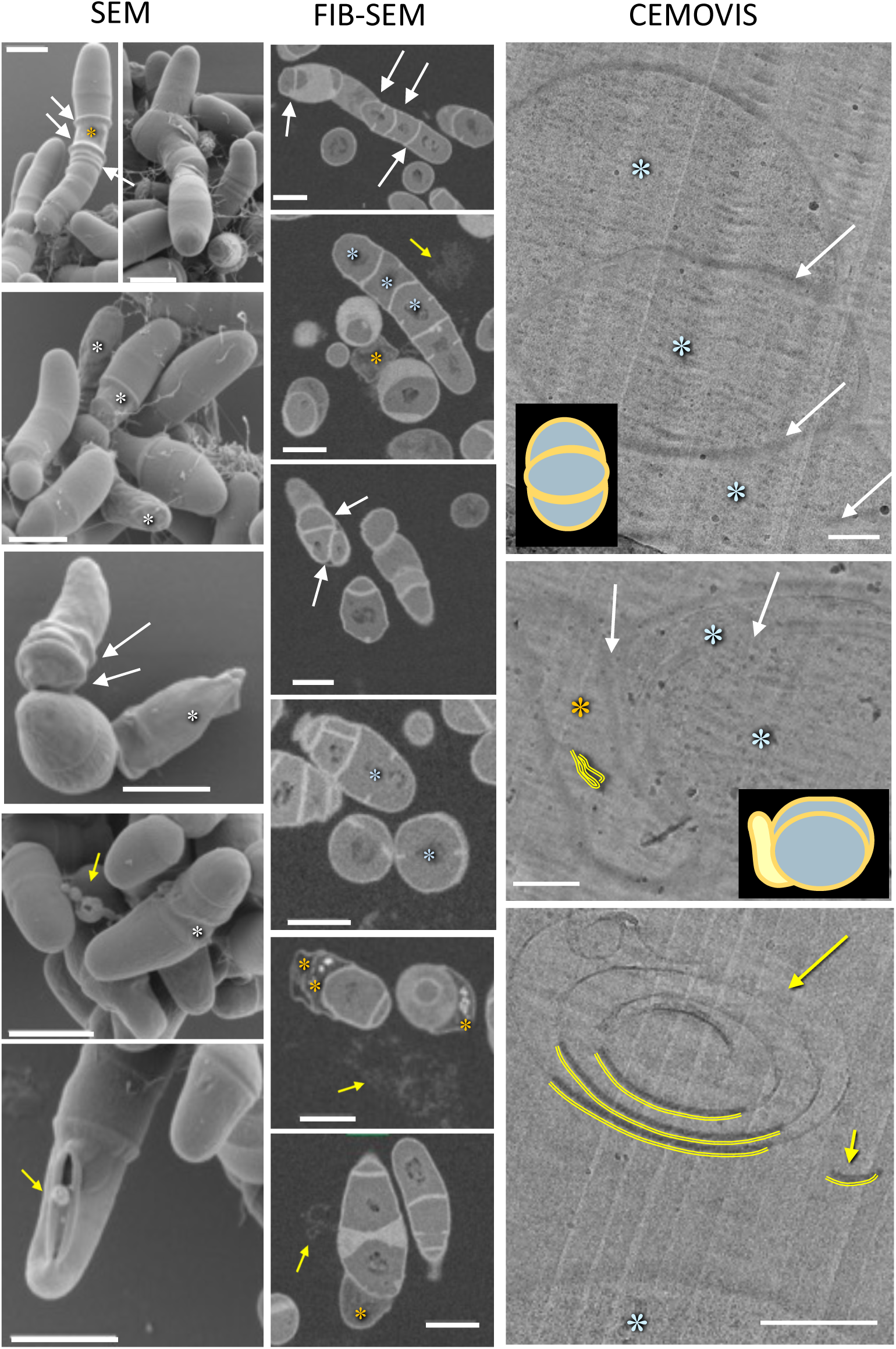
Electron microscopy observations of Δ*myts* cells. **(SEM)** Multiple septa are visible (white arrows), along with deformed cellular compartments (white asterisks). (**FIB-SEM**) Multiple septa separate distinct compartments that are either filled with cytoplasm (blue asterisks) or appear empty/clear (orange asterisks). Extracellular membrane structures are indicated by yellow arrows. (**CEMOVIS**) Overview images reveal a series of septa dividing the cell into compartments that are either cytoplasm-filled (blue asterisks) or quasi-empty (orange asterisks), containing cytoplasmic remnants and membrane debris (underlined in yellow). Extracellular multilayered membrane structures (vesicles, debris) are also observed (bottom panel, yellow arrows, locally underlined in yellow). Scale bars 1 µm (SEM and FIB-SEM), 200 nm (CEMOVIS).

To relate these structural defects to division dynamics, we monitored Δ*myts* growth by time-lapse microscopy alongside WT cells. As expected, WT cells undergo V-snapping division (Fig. 5 and Video 1), the canonical mode of separation in Mycobacteriales, including *Mycobacterium tuberculosis*. In contrast, Δ*myts* cells elongate extensively (Fig. 5 and Video 2) and seems to accumulate multiple septa that subdivide the cytoplasm into discrete compartments, similarly to what is observed on standard DIC and Van-FL labeled cells in Fig. S7. In some cases, division appears to occur through the classical snap event, whereas in other cases, cells undergo through a lysis event occurring between two compartments that remain viable (Fig. 5, red arrow and Video 2). We also observed that snapping appears more readily when cells are in contact with neighbors, suggesting that mechanical pressure exerted by adjacent cells promotes breakage, a phenomenon previously noted in the literature [29].

**Figure 5:**
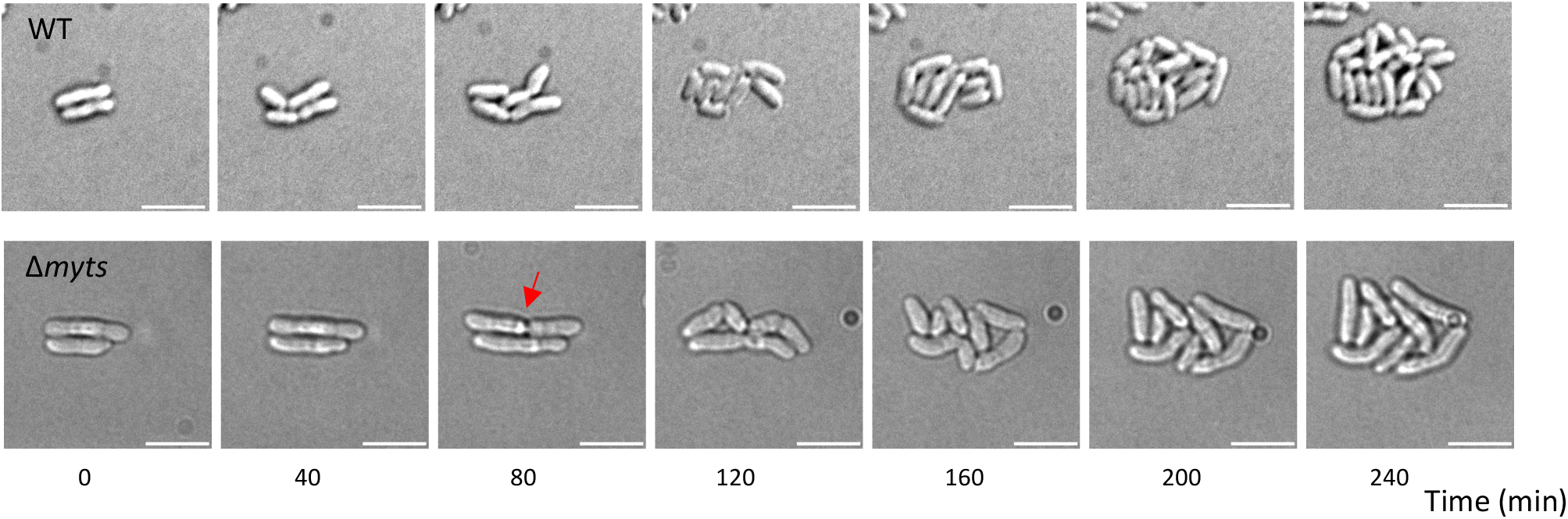
Time-lapse microscopy observation of WT and Δ*myts* cells. Images are recorded at 20-min intervals (see Video). DIC images are presented at the indicated time point (40 min intervals). The red arrow indicated a depression zone in the bacterial cell, consistent with lysis, of part of the cells. Scale bar, 5 μm. Videos are available in electronic supplementary data (video 1 for WT and video 2 for Δ*myts*).

### Abnormal morphology of Δ*myts* mutant is not explained by secondary mutations

Given that the Δ*myts* and Δ*pks* mutants both lack a mycomembrane but exhibit distinct morphological phenotypes, we investigated whether secondary mutations could account for the abnormal morphology of the Δ*myts* mutant. Whole-genome sequencing of five independent Δ*myt*s clones confirmed the deletion of all six *myt* genes and revealed a small number of secondary nucleotide substitutions relative to the WT strain. As detailed in Table S6, some mutations were clone-specific and therefore unlikely to account for the shared abnormal morphology observed in all five Δ*myt*s clones (Fig. S15). Other mutations were shared across multiple clones and may have arisen early during the deletion procedure. These included synonymous and intergenic substitutions, as well as a mutation in the upstream region of the *mscL* gene (encoding a high-conductance mechanosensitive ion channel). MscL is an inner membrane channel involved in the response to osmotic stress and is not expected to alter cell morphology or cell division [30] Nevertheless, a contribution of these secondary mutations to the observed phenotype cannot be completely excluded. Based on our previous observations that individual deletion of *mytA*, *mytB*, and *mytC* affected cell morphology and/or aggregation to varying degrees, we performed complementation experiments with these three genes to further investigate their contributions to the Δ*myts* phenotype and determine whether the observed defects could be partly attributable to polar effects associated with the sequential deletions. Complementation with *mytA* substantially restored the WT-like phenotype, whereas complementation *mytB* led to a more partial restoration, with only a subset of cells recovering a WT-like phenotype. In contrast, no apparent phenotypic restoration was observed upon complementation with *mytC* (Figure S16). These results are consistent with our observations of the individual deletion mutants and support a major contribution of MytA, and a lesser contribution of MytB, to maintain normal cell morphology. Although no apparent restoration was observed upon complementation with *mytC*, an additional contribution of MytC, as well as of the other mycoloyltransferases (MytD, MytE, and MytF), cannot be excluded. Thus, the pronounced morphological defects of the Δ*myts* mutant may result from the combined loss of multiple mycoloyltransferases, with MytA and MytB playing particularly important roles.

### Abnormal morphology of Δ*myts* mutant is not suppressed by *pks* deletion

We next sought to determine whether the accumulation of TMM, absent in the Δ*pks* strain, could account for the morphological divergence observed between the strains. To test the hypothesis that TMM accumulation in the Δ*myts* mutant contributes to the observed division defect, we deleted the *pks* gene in the sextuple mutant background and examined whether this genetic modification rescued the elongated and multisepta phenotype. The Δ*myts*Δ*pks* resulting mutant, was confirmed by PCR to lack the *myt* genes and no longer synthesizes TMM or TDM (Fig. S17A), similar to the Δ*pks* strain. Whole-genome sequencing of three Δ*myts* Δ*pks* clones confirmed the expected deletions and revealed very few clone-specific additional nucleotide substitutions (Table S7), including mutations located upstream of the pks gene, as well as missense mutations. All three clones also shared a deletion in a gene encoding a hypothetical protein, resulting in the loss of a single isoleucine residue within a stretch of four consecutive isoleucine residues in the protein sequence. Critically, the abolition of mycolate production does not alter the phenotype associated with the Δ*myts* mutations. All Δ*myts*Δ*pks* mutant clones retain similar severe cell division defects and elongated cell morphology characteristics of the original Δ*myts* mutant, as shown by differential interference contrast (DIC) microscopy (Fig. S15). Notably, as observed by scanning electron microscopy, this strain exhibits a clear multiseptal phenotype accompanied by an elongated morphology very similar to the one observed in the Δ*myts* strain and clearly different from the one observed in the Δ*pks* (Fig. 6). Although the morphology of these mutants is complex and heterogeneous, we performed a global flow cytometry to evaluate relative cell size (forward scatter, FSC) and granularity (side scatter, SSC) across a large population (50,000 cells) (Fig. 6C and Fig. S17B-S17C). Scatter plots of FSC versus SSC from four independent exponential-phase cultures revealed that the Δ*myts*Δ*pks* mutant clusters closely with the Δ*myts* strain and remains clearly distinct from the *Δpks* and WT strains. Altogether, these results support the conclusion that the division defect is not rescued by the loss of mycolate synthesis and is consistent with the Δ*myts* phenotype being epistatic to the Δ*pks* phenotype.

**Figure 6:**
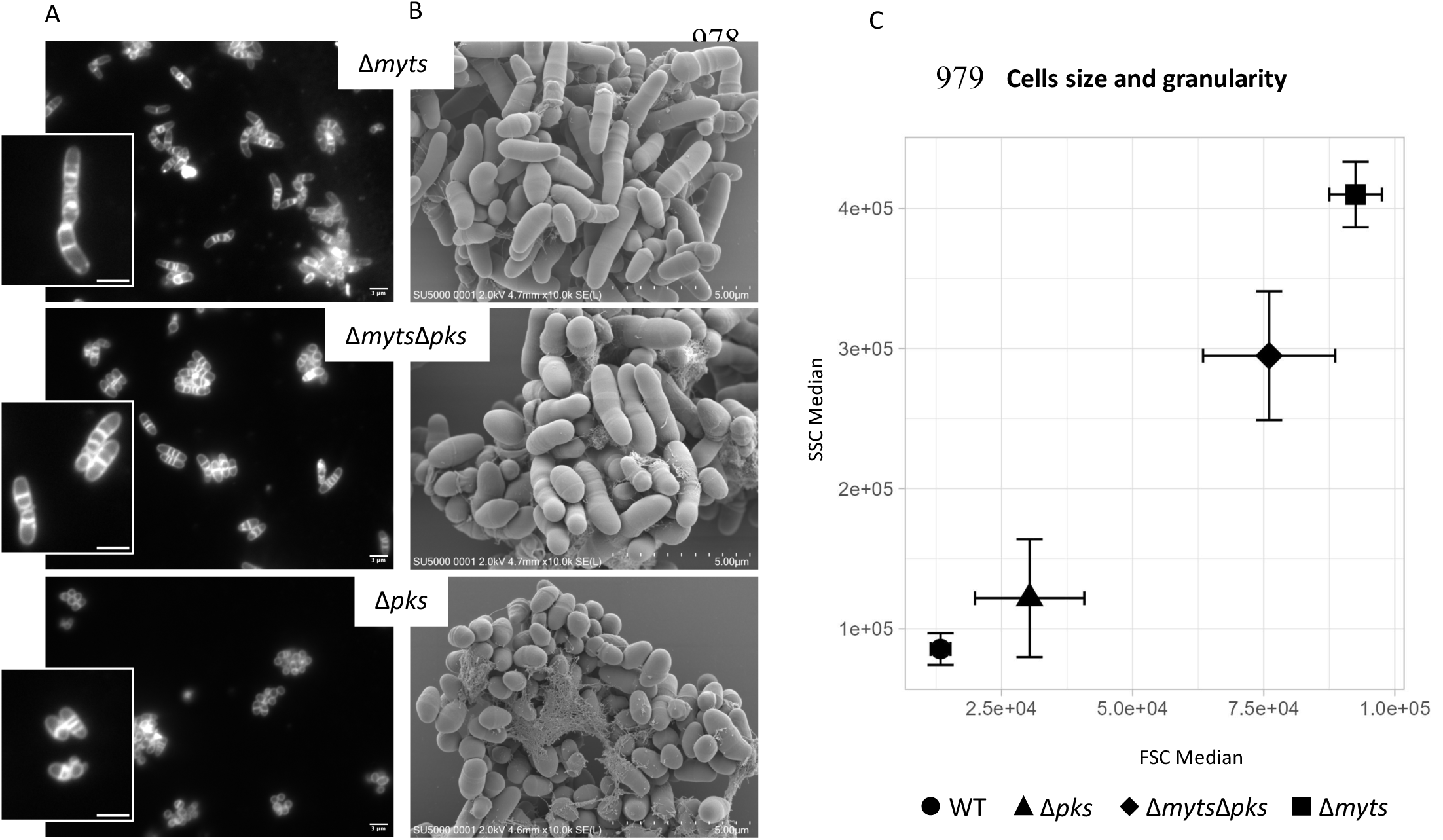
Morphology of Δ*myts*, Δ*pks* and Δ*myts*Δ*pks* mutants. **(A)** Fluorescent labeling of membranes by Nile red (NR): The Δ*myts*Δ*pks* mutant displays elongated cells containing multiple septa, resembling the Δ*myts* phenotype. Daughter cells are frequently observed trapped wlabeled at the cell periphery and only occasionally exhibits one or two-three septa. Zoom of representative cells in the squares. Scale bar, 3 μm. **(B)** SEM micrographs: in contrast to the coccoid and stocky ithin the mother cell, as also seen in Δ*myts*. In contrast, the Δ*pks* strain is primarily Δ*pks*, Δ*myts*Δ*pks* is much more elongated and irregularly shaped as observed for the Δ*myts*. Scale bar, 5 μm. **(C)** Scatter plot of flow cytometry data showing the median SSC value as a function of the median FSC value for 4 independent biological replicates and 2 technical replicates each (the corresponding box plots are shown in **Figure S17B**).

## DISCUSSION

### All Myts are dispensable for viability but contribute to normal cell morphology

The construction of the Δ*myts* sextuple mutant demonstrates that all six mycoloyltransferases can be simultaneously deleted in *C. glutamicum* without abolishing viability. However, this genetic background is associated with pronounced growth and morphological defects. Whole-genome sequencing of five independently obtained Δ*myts* clones revealed distinct sets of secondary mutations (except a mutation upstream *mscL*), making it unlikely that a single secondary mutation accounts for the morphological defects shared by all clones. Nevertheless, mutations affecting the RsdA–σᴰ regulatory pathway were identified in two independent clones, one in *rsdA* and the other in *sigD*. The occurrence of mutations affecting two components of the same regulatory pathway in independent clones is noteworthy and may reflect convergent adaptation. As the RsdA–σᴰ system regulates several cell-envelope-associated functions, including mycolic acid metabolism, these mutations could alter σᴰ-dependent gene expression and potentially modulate mycolic acid/TMM synthesis [32], [33]. Such changes might partially alleviate the lipid imbalance resulting from the absence of the Myts enzymes and thereby improve the fitness of the Δ*myts* mutant. However, this hypothesis remains speculative and requires further investigation.

The individual deletion mutants and complementation experiments both support distinct contributions of individual mycoloyltransferases to cell morphology, particularly MytA and, to a lesser extent, MytB. The elongated morphology of Δ*mytA* cells and the rounder shape of Δ*mytB* cells point to distinct roles for these enzymes in maintaining normal cell morphology. Consistent with this, complementation with *mytA* substantially restores a wild-type-like phenotype in the Δ*myts* background, whereas *mytB* complementation provides only partial restoration. The substantial rescue by *mytA* also argues against the morphological defects being explained solely by polar effects arising during the sequential deletion procedure. The contribution of MytC is less clear, as Δ*mytC* cells display aggregation and occasional elongation, but no apparent morphological rescue was observed upon complementation. Nevertheless, the relatively mild phenotype of Δ*mytC* suggests that the severe division defects of Δ*myts* cannot be explained simply by the loss of protein mycoloylation. This interpretation is consistent with published O-mycoloylome datasets, in which proteins with established roles in cell division are not significantly represented among the identified substrates. Although RipA has been proposed as a potential candidate in one study [34], its O-mycoloylation was not experimentally demonstrated by the authors, and no functional role for this putative modification in cell division has been established. Importantly, beyond the individual contributions of MytA, MytB and MytC, the progressive emergence of morphological defects in the intermediate mutants indicates that the contributions of the other Myts cannot be excluded. Since the genes were deleted sequentially, the order of deletion and potential interactions between the resulting mutations may also influence the observed phenotypes. Thus, the severe defects of Δ*myts* likely reflect the combined loss of multiple mycoloyltransferases and their contributions to cell-envelope organization and cell division.

### Arabinogalactan-bound mycolates are essential for initiating mycomembrane assembly

Although it has long been established that mycolic acid chains are synthesized in the cytoplasm by polyketide synthases, the origin of TMM itself remains unclear. In *C. glutamicum*, TMM formation was initially proposed to occur in the cell envelope via a mycoloyltransferase that would transfer cytoplasm-derived mycolic acids, after their translocation across the inner membrane, onto trehalose [35], [36], [37]. Later, *in vitro* biochemical studies on Pks13 from *M. tuberculosis* [3] suggested that this enzyme could in fact directly transfer mycolic intermediates onto trehalose to form a cytoplasmic TMM precursor, subsequently exported across the inner membrane. In the present study, we show *in vivo* that TMM is still synthesized in the *C. glutamicum* Δ*myts* mutant, providing direct physiological evidence that TMM formation clearly occurs independently of mycoloyltransferases and supporting the model in which TMM is synthesized in the cytoplasm by polyketide synthases in both organisms. As a consequence, the Δ*myts* mutant constructed in this study is a powerful system to uncouple early and late steps of mycolic acid metabolism and directly assess the contribution of downstream transfer reactions to mycomembrane biogenesis. Interestingly, despite producing large amounts of TMM, the Δ*myts* mutant likely continues to flip TMM across the inner membrane, as suggested by the detection of TMM-containing vesicles in the culture supernatant. This is consistent with previous work showing that MmpL3 mediates the flipping of TMM to the outer leaflet of the inner membrane in *M. smegmatis* spheroplasts [38]. Nevertheless, the Δ*myts* mutant fails to assemble a mycomembrane, indicating that TMM alone is insufficient to support its formation and suggesting that additional mycolate-containing components are required. Current evidence demonstrates that TDM is not essential in *Mycobacterium smegmatis*: a recently characterized TDM-deficient mutant fails to synthesize TDM yet retains the ability to produce arabinogalactan-linked mycolates and exhibits normal growth [39]. We therefore propose that the lack of a mycomembrane in the Δ*myts* is due to the absence of arabinogalactan-linked mycolates, which likely represent the critical determinant of mycomembrane biogenesis. In their absence, the outer membrane may either fail to assemble properly or become destabilized, thereby promoting the release of TMM-containing vesicles into the culture supernatant, as observed in our mutant. This is also consistent with our previous observations in the Δ*aftB* mutant, where reduced arabinogalactan mycoloylation results in a destabilized and partially detached mycomembrane. [22]. An additional, non-mutually exclusive possibility is that the inability to transfer mycolates from TMM onto arabinogalactan leads to the accumulation of TMM in the outer leaflet of the inner membrane. Such an accumulation could alter the biophysical properties of the inner membrane and compromise its stability, thereby contributing to vesicle formation and release. This hypothesis is consistent with the detection of small amounts of inner membrane-derived lipids (PIMs and PI) among the vesicles secreted by the Δ*myts* mutant. From a broader perspective, this situation is reminiscent of lipopolysaccharide-containing diderm bacteria, in which the outer membrane is tethered to the cell wall by OmpA and the Braun lipoprotein (Lpp), two major structural components that are required for maintaining envelope integrity [40]. By analogy, arabinogalactan-linked mycolates may play a comparable anchoring role in mycolic acid-containing diderm bacteria, where such coupling could be even more critical and may constitute a prerequisite for initiating the assembly of the outer membrane. This increased dependence may reflect fundamental differences in cell envelope organization and protein equipment, particularly during septation. In LPS-diderm bacteria such as *Escherichia coli*, the Tol-Pal system couples the inner and outer membranes and coordinates their constriction during division [41]. In Mycobacteriales, which lack this system, covalent attachment of mycolates to arabinogalactan ensures strong outer membrane attachment to the cell wall, a likely critical requirement for coordinated envelope assembly during septation.

### Mycoloyltransferases, not their metabolites, play a critical role in late cytokinesis

Although both Δ*myts* and *Δpks* lack a detectable mycomembrane, their phenotypes differ strikingly. Δ*pks* cells remain relatively homogeneous and do not display major septation defects, whereas Δ*myts* cells are highly elongated, multiseptated, and morphologically heterogeneous. Importantly, deletion of *pks* in the Δ*myts* background does not suppress this phenotype, despite eliminating TMM accumulation. These findings demonstrate that the division defect is independent of mycolic acid synthesis, TMM buildup, or simple loss of the mycomembrane. Loss of mycoloyltransferases is therefore not phenotypically equivalent to loss of mycolic acids and blocking the transfer step produces cellular consequences that cannot be explained solely by absence of the end products or the accumulation of intermediates, i.e. TMM as shown by the still defective morphology of the Δ*myts*/Δ*pks* mutant. This observation is conceptually important as mycolate synthesis and transfer are often considered parts of a linear pathway [31], [36].

Elongated morphology in bacterial mutants is frequently associated with division defects. Such phenotypes may arise either from impaired septum formation (e.g., defective Z-ring assembly) or from failure of daughter-cell separation after septation. In Δ*myts*, multiple lines of evidence indicate that septum formation occurs: septa are labeled by vancomycin and Nile Red, and are clearly visible by CEMOVIS and FIB-SEM. Thus, divisome assembly and septal peptidoglycan synthesis are functional. The defect therefore most likely lies in late stages of division, specifically septal maturation or daughter-cell separation. Several electron microscopy approaches and time-lapse microscopy show that Δ*myts* cells initiate septation but accumulate multiple septa that subdivide the cytoplasm into compartments. In fact, the envelope ruptures and vesicle release observed in Δ*myts* are not merely the result of envelope instability in the absence of a mycomembrane, as Δ*pks* cells do not display comparable fragility. The curved, rounded, or closely spaced septa observed ultrastructurally suggest defective coordination between septal peptidoglycan synthesis and septal resolution.

### A possible regulation of septal hydrolases by Mycoloyltransferases ?

In Mycobacteriales, daughter-cell separation depends on tightly regulated peptidoglycan hydrolases that remodel the septum prior to mechanical snapping [25], [42]. In *Mycobacterium tuberculosis*, the endopeptidase RipA plays a central role in septal cleavage [43], [44], [45], while in *Corynebacterium glutamicum*, additional enzymes such as Cg2402 might also contribute to septal processing and efficient cell separation [46]. Loss of these hydrolases leads to chaining, multiseptation, and “bamboo-like” morphologies with rounded, unresolved septa; phenotypes that closely resemble those observed in Δ*myts* mutants. These striking similarities raise the possibility that mycoloyltransferases influence septal hydrolase function. Because both enzyme classes operate within the multilayered cell envelope, Myts could participate in coordinating outer envelope remodeling with septal peptidoglycan cleavage. In this speculative view, Myts might act as spatial regulators, scaffolding components, or modulators of local envelope architecture, thereby creating a permissive environment for hydrolase access or activation at division sites. In their absence, septa may assemble but fail to mature or become mechanically competent for splitting, leading to repeated rounds of cytokinesis without effective daughter-cell separation.

The absence of comparable division defects in Δ*pks* cells argues against a mechanism strictly dependent on mycolic acid transfer itself. Instead, this observation supports the possibility of a moonlighting function for Myts independent of their canonical catalytic activity, or a structural role in organizing envelope layers during septation. Interestingly, MytA has been previously suggested to make a physical link between the mycomembrane and the underlying peptidoglycan-arabinogalactan polymer that is essential for envelope integrity as assessed by antibiotic susceptibility [47]. Alternatively, Myts may retain the ability, even in a Δ*pks* background, to transfer conventional fatty acids onto trehalose or arabinogalactan. However, such residual activity would be expected to be minimal given that it is not sufficient to support the formation of a bona fide mycomembrane in the Δ*pks* strain. Under this scenario, it would not be the presence of a mature mycomembrane per se, but rather subtle lipid modifications or local envelope properties that modulate septal hydrolase activity. Discriminating between these models remains challenging. As a result, it is currently unclear whether Myts primarily affect hydrolase localization, activation, or instead the physical properties of the septal envelope itself.

### Concluding remarks and perspectives

By genetically uncoupling mycolate synthesis from mycolate transfer, our work reveals that mycoloyltransferases play two mechanistically distinct and functionally critical roles in *C. glutamicum*. First, we demonstrate that the transfer of mycolates onto arabinogalactan by mycoloyltransferases is a decisive step in initiating mycomembrane assembly. Although this function is consistent with their annotated enzymatic activity, our data provide direct genetic evidence that mycolate transfer on AG, rather than mycolate synthesis per se, is the key determinant for triggering higher-order envelope organization. Second, and unexpectedly, we uncover a role for mycoloyltransferases in late cytokinetic events: these enzymes are required for proper septal maturation and efficient daughter-cell separation independently of mycolic acid production itself. Together, these findings reposition mycoloyltransferases as central coordinators of both envelope biogenesis and division fidelity.

The Δ*myts* phenotype reveals a striking specificity for late steps of cytokinesis, particularly septal resolution, highlighting the specific requirement for mycoloyltransferases during these stages. This suggests that outer envelope remodeling must be tightly coordinated with septal peptidoglycan hydrolysis and the mechanical forces driving V-snapping division. One parsimonious interpretation is that mycoloyltransferases help synchronize envelope maturation with septal cleavage, ensuring that mechanical separation occurs only once the envelope has reached a permissive structural state. In this framework, they could contribute to a functional checkpoint linking envelope assembly to daughter-cell separation. Mechanistically, several non-mutually exclusive scenarios can be envisioned. Mycoloyltransferases may exert structural roles at division sites, modulate septal hydrolase activity by shaping the local envelope environment, or integrate into divisome-associated protein networks that coordinate multilayered envelope remodeling. Clarifying these possibilities will require defining their spatial organization during division and identifying potential interaction partners. Because polar growth and V-snapping division are conserved across Mycobacteriales [48], [49], it is tempting to speculate that such coupling between envelope remodeling and late cytokinesis may represent a broader principle within this order. Testing this hypothesis in other species, including pathogenic mycobacteria, will be necessary to determine whether the dual role of mycoloyltransferases in envelope biogenesis and cytokinesis is evolutionarily conserved. More broadly, our findings underscore that enzymes classically assigned to lipid biosynthetic pathways can exert regulatory control over cell morphogenesis and division. Disrupting mycoloyltransferase function may therefore compromise not only envelope architecture but also the coordination and robustness of bacterial cytokinesis, offering new conceptual perspectives on envelope-centered vulnerabilities in Mycobacteriales.

## MATERIALS & METHODS

### Bacterial strains, plasmids and culture conditions

Strains and plasmids used in this study are listed in de Table 1. Cultures were performed in Luria Bertani (LB) medium at 37°C for *E. coli* and in Brain Heart Infusion (BHI) medium at 25 or 30°C for *C. glutamicum*. When appropriate, kanamycin (Km) is added to a final concentration of 25 µg/ml and chloramphenicol (Cm) at concentrations of 30 µg/ml or 6µg/ml respectively for *E. coli* and *C. glutamicum*. The liquid cultures were grown under agitation (190rpm) and monitored by measuring the optical density at 600nm (OD_600nm_).

### Plasmids and strains construction

Oligonucleotides used in this study are given in the Table S1 (supplementary data). The Δ*myts* mutant strain of *C. glutamicum* was constructed using the technique described by Schäfer *et al*. [21], which consists in deleting a gene without leaving any resistance gene cassette marker, thus enabling successive gene deletions. Briefly, a 400bp fragment upstream of the gene and a 400bp fragment downstream of the same gene are amplified by PCR respectively with the del1/del2 and del3/del4 gene corresponding oligonucleotides. These fragments are cloned in juxtaposition in the pk18mobsac vector. The corresponding plasmid pk18del obtained in *E. coli* DH5α is sequenced by the respective ver1 and ver2 oligonucleotides for verification. The plasmid obtained is then electroporated into ATCC13032 res-competent cells [50]. The following steps described by Shäfer *et al.* with isolation in BHI-10% sucrose and screening of Km sensible colonies allows the selection of genomic deletion mutants of the gene of interest by PCR with the respective ver3/ver4 oligonucleotides.

The PCR reactions were carried out using DreamTaq or Phusion™ High Fidelity DNA polymerases from ThermoFisher Scientific, and DNA purification was performed using Nucleospin kits for gel and PCR purification or plasmid purification from Macherey-Nagel. Oligonucleotide synthesis and DNA sequencing were performed by Eurofins.

The *myt*B ORF was amplified by PCR with primers *myt*B-fw and *myt*B-rv, digested with HindIII, and ligated into a HindIII/EcoRI(blunt) digested pXMJ19, transformed into *E. coli*DH5α, and recombinant clones were screened by restriction analysis and sequencing.

### Genome sequencing

Genomic DNA were extract with the Nucleospin Microbial DNA kit from Macherey-Nagel followed by ethanol precipitation. Genomic DNA libraries were then prepared with NEB reagents (NEBNext DNA Fragmentase M0348S, Ultra II End Repair/dA-Tailing Module E7546S and quick ligation module E6056S). The quality of the final libraries was assessed with an Agilent Bioanalyzer, using an Agilent High Sensitivity DNA Kit. Libraries were pooled in equimolar proportions and sequenced using paired-end 2×75 pb runs, on an Illumina NextSeq550 instrument, using NextSeq 550 High 150 cycles kit, or with paired-end 2×61 pb reads, on an Illumina NextSeq2000 instrument, using a NextSeq 2000 XLEAP P2 100 cycles kit. Demultiplexing has been done (bcl2fastq2 V2.18.12 for Nextseq550 and bcl-convert 4.4.6 for NextSeq2000) and adapters removed (Cutadapt1.15 for NextSeq550 and Cutadapt 5.2 for NextSeq2000), only reads longer than 10pb were kept for analysis. Sequenced reads (read 1 only) were mapped with bwa (0.7.17-r1188) to access NC_006958. The mapped reads were used as input in Freebayes (v1.0.2-16-gd466dde) to detect SNPs. SNPs common with those present in the reference sample (WT for Δ*myts* variants and Δ*myts* for Δ*myts*Δ*pks* variants) were removed for each sample with VarScan (v2.3.9). Variation annotation was performed with SNPeff (v4.3t) using the *Corynebacterium glutamicum* ATCC 13032 database. Only variations with an allele frequency of 100% and a depth greater than 10 reads were selected. Paired-end bwa mapping was used with Breakdancer (breakdancer-max1.4.5-unstable-66-4e44b43) to detect larger structural rearrangements. The results of the comparison between the Δ*myts* variants and the WT strain are presented in Table S6, while the comparison between the Δ*myts* and Δ*myts*Δ*pks* variants is reported in Table S7.

### Differential interference contrast and fluorescence microscopy

Live cell imaging was carried out spotting a culture sample (in exponential or stationary phase) in 1% agarose pad glass slide and covered by a coverslip before imaging.

Nile red or NR (9-(Diethylamino)-5*H*-benzo[*a*]phenoxazin-5-one), rDHPE (rhodamine B 1,2-dihexadecanoyl-sn-glycero-3-phosphoethanolamine, triethylammonium salt) and BODIPY FL-conjugated vancomycin (Van-FL) from Invitrogen were used to label respectively membrane, mycomembrane and peptidoglycan. The cells were harvested and washed twice with phosphate buffered saline (PBS 1x). For 100µL of cell suspension, 2 or 1µL of 100µg/ml stock solution of NR, van-FL, or rDHPE fluorescent dyes were added for 20 min of staining at 25°C. The cells were then washed 3 times (except for NR staining) before being deposited on a glass slide coated with an agarose (1%) layer and covered by a coverslip.

During time-lapse, cells were grown in BHI. A fresh culture at an initial OD_600nm_ between 0,2 and 0,4 were grown further in shaking flasks for 1h-1h30 at 30°C, 190 rpm. The exponentially growing cells were then spread onto BHI agar pad slide glass, and imaged at twenty-minute intervals.

Images were taken on a three-dimensional deconvolution microscope (DMIRE2, Leica Microsystems) coupled to a 20-MHz Cool SNAP-HQ2 CCD camera (Roper Technologies) using a HCxPL APO 100× oil CS objective, NA = 1.40 (Leica Microsystems) and a Lumencor LED system (Optoprim) (485/20nm with a GFP filter for viewing FL-Van, 586/20nm with mCherry filter for viewing r-DHPE or NR). A total of 16 Z-stack images, each spaced 0,2 µm, were acquired and deconvoluted with MetaMorph software (Molecular Devices). The images obtained were processed using Metamorph or FIJI.

### Lipid analysis

#### Purification of trehalose mycolates

As a general rule, 20 mL of culture were harvested during the stationary phase (after 24 h of growth at 30 °C). Lipids were extracted from the resulting wet cell pellets by overnight incubation at room temperature in 6 mL of a CHCl₃/CH₃OH mixture (1:2, v/v). Following this first extraction, the delipidated cell pellets were retained for subsequent analysis of cell wall-bound mycolates (see below). The organic phase was then evaporated to dryness, and the extracted lipids were resuspended in 100 µL of CHCl₃. Lipid analysis was performed by thin-layer chromatography (TLC) on silica gel 60 plates (Macherey-Nagel) using CHCl₃/CH₃OH/H₂O (65:25:4, v/v/v) as the developing solvent. Lipids were visualized by immersing the TLC plates in 10% H₂SO₄ in ethanol, followed by heating at 110 °C. The lipid classes were identified by their Rf and the precise molecular identities of TMM and TDM were further confirmed by mass spectrometry.

#### Purification of cell wall-bound mycolates

Delipidated cell pellets (see above) were subjected to methanolysis with 3 M HCl in CH₃OH to generate mycolic acid methyl esters (MAMEs). The resulting MAMEs were recovered by successive liquid–liquid extractions with CHCl₃/H₂O (1:1, v/v). The combined organic extracts were analyzed by thin-layer chromatography (TLC) on silica gel 60 F254 plates (Merck). They were separated by developing the plates twice at 4 °C in hexane/ethyl acetate (95:5, v/v). MAMEs were visualized by spraying the plates with phosphomolybdic acid in ethanol, followed by charring. The identities of the mycolates were confirmed by mass spectrometry.

#### Purification of secreted mycolates

Vesicles were purified from the supernatant of a 100 mL BHI culture. Cells were removed by centrifugation at 4,000 × *g*, and the clarified supernatant was subsequently centrifuged at 35,000 × *g* for 2 h using a 45Ti rotor to pellet vesicles. The pellet was resuspended in 25 mM HEPES buffer (pH 7.5) and layered onto a discontinuous sucrose cushion consisting of 20% (w/w) sucrose over a 60% (w/w) sucrose cushion. After centrifugation for 3 h in a TLA100.2 rotor at 32,000 × *g*, the vesicles collected at the 20%/60% interface were recovered, diluted in 50 mL of 25 mM HEPES buffer (pH 7.5), and centrifuged again at 35,000 × *g* for 2 h in a 45Ti rotor. The final pellet was resuspended in HEPES buffer and stored at −20 °C.

#### Analysis of trehalose mycolates by MALDI and NMR

TMM and TDM were analyzed by Matrix-Assisted Laser Desorption and Ionisation coupled to Quadrupole Ion Trap Time Of Flight (MALDI Q-IT TOF) MS analysis. Samples dissolved in CHCl_3_/CH_3_OH (2:1) were mixed (1:1) with 10 mg/mL 2,5-dihydroxy benzoic acid (DHB) in CHCl_3_/CH_3_OH (1:2) spotted on the target before analysis on a Axima Resonance under Shimadzu Biotech MALDI-MS (SHIMADZU, Kyoto, Japan). Laser energy was optimized for each sample between 90 and 130 eV, and acquisition was conducted in positive reflectron mode.

NMR spectra were acquired at 293 K TBI probes and AVANCE Neo consoles (BRUKER BIOSPIN GmbH, Rheinstetten, Germany), with resonance frequencies of ^1^H at 400 MHz on samples solubilized in CDCl_3_/CD_3_OH (2:1). Data were analyzed with Topspin 4.0.6 software.

#### CEMOVIS

Bacteria cultured on agar Petri dishes were harvested on the tip of a pipette, gently mixed with and equal volume of dextran (MW 40kD, Euromedex) 40 % in PBS, and immediately deposited into a gold/copper cupel for vitrification in an EMPACT2 high pressure freezer (Leica). Cupels were mounted into the flat specimen holder of a cryo-ultramicrotome FC6/UC6 (Leica) cooled down to −140°C. After trimming of a ∼ 50 x75 mm pyramid, 2 to 3 mm-long ribbons of cryo-sections were obtained with a 25° diamond knife (Diatome), using a double micromanipulator (Leica) [51] at a cutting speed of 0.4 mm/sec and a nominal feed of 40 or 50 nm. Ribbons were collected and sticked onto Quantifoil S7/2 grids covered with an ultrathin (1-2 nm) continuous carbon film using a Crion electrostatic Gun [52].

Grids were monted in a Gatan 626 cryoholder and observed in a JEOL 2O10F operated at 200kV. Images were acquired at 2000 or 3000 nm defocus with a Gatan K2 summit camera, at pixel sizes of 0.3808729, 0.4411774, or 0.6233850 nm.

### Cryo-Transmission Electron Microscopy (Cryo-TEM)

A 4 µL aliquot of sample was adsorbed onto a glow-discharged (Qorum, UK) holey carbon-coated grid (Lacey, Tedpella, USA), blotted 4 seconds with Whatman 1 filter paper and vitrified into liquid ethane at −180 °C using a Leica GP2 plunger (Leica microsystems, Austria). Frozen grids were transferred onto a Talos 200C electron microscope (FEI, USA) operating at 200KV using a Gatan 626 side-entry cryoholder. Micrographs were collected on a Ceta CMOS camera at a pixel size of 0.2 nm.

### SEM

Culture samples of exponential phase (4-6 h of growth) or stationary phase (24h of growth) were fixed with Glutaraldehyde 2% in Na-Cacodylate buffer 0.1 M (pH 7.4) 2h at RT and further ON at 4°C. For that, the samples were deposited on 1cm square microscopy glass slide, cut with a diamond pen. Glass slides were previously cleaned with 70% ethanol, 5 min plasma cleaner, 10 min sonication bath in ethanol 100%, and in MilliQ water, and finally coated with 0,1M polylysine (Merck Sigma Aldrich) during 5 min, then rinsed with MilliQ water.

Samples were rinsed twice during 10 min in 0.2 M sodium cacodylate solution (pH 7.4) and underwent progressive dehydration by soaking in a series of ethanol (50%, 70%, 90%, 100%, anhydrous 100%) before critical-point drying under CO2 (Leica EM300 CPD, slow 20 exchange cycles, 2min delay). Samples were mounted on aluminum stubs (15 mm and 32mm diameter) with carbon adhesive discs (LFG France), and were coated with Au/Pd (Quorum SC7620, 5 Pa of Ar, 270 seconds of sputtering at 2.5 mA). Samples were then observed in high-vacuum by scanning electron microscopy (FEG-SEM-LV SU-5000, Hitachi, Milexia, 91190-Saint-Aubin, France), at 2KeV and 30 spot size, SE(L) detector, and 5mm WD.

### FIB-SEM

Bacteria aggregates, from stationary phase of 24 h growing cells, were fixed 2h at RT in 2.5 % glutaraldehyde in 0.1 M phosphate buffer at pH 7.4, then rinsed in buffer and post-fixed in 1 % osmium and 1.5 % potassium ferrocyanide for 1 h at RT. Samples were then rinsed in water and dehydrated in graded acetone series (30-50-70-90-100-100 %, 10 min per bath) and embedded in araldite resin with increasing concentration of resin mixed with acetone (30 %-50 %-70 %) for 2 h in each bath, and overnight in pure araldite. Sample were then incubated 72 h at 60 °C for resin polymerisation.

The block was cut parallelly to its surface in order to be 2–3 mm high, and mounted on an SEM stub (Agar Scientific) using silver paint (Agar Scientific, ref AGG3691). Silver paint was further added around the block surface. The sample underwent platinum sputtering of 20 nm (Safematic CCU-010) before insertion in the FIB-SEM chamber. FIB-SEM imaging was performed using a Zeiss Crossbeam 550, following the Atlas 3D nanotomography workflow. FIB milling was performed at 700 pA. SEM imaging was done with an acceleration voltage of 1.4 kV and a current of 400 pA using an energy-selective backscattered (ESB) detector (ESB grid 1000 V). Imaging was done using a 25 nm isotropic voxel size. Segmentation was generated using Microscopy Image Browser (MIB) and 3D visualization and analysis performed with Dragonfly software (non-commercial licence).

### Length shape measurement

To measure the size of bacteria, and given the difficulty in obtaining individual cells for conventional phase contrast segmentation, we used AI to assist us in a semi-manual DIC segmentation under conditions where cells are least aggregated (25°C). Masks Segmentations were generated using Microscopy Image Browser (MIB) and analyzed by MicrobeJ plugin in FIJI. GraphPad Prism 10 software was used to generate Superplots and performed statistical analyses. Differences were considered significant when the calculated *p*-values were smaller than 0,05.

### Cytometry

A fresh culture was prepared to an initial OD_600nm_ 0,1-0,3. After 4 hours of growth at 30°C (190rpm), 2-4 ml of cultures were harvested, washed and resuspended in 1 ml of PBS at approximately OD_600nm_ 0,25 for flow Cytometry analysis. 50,000 cells per replicate were detected with a speed of 10µL/min on a Cytoflex S bench-top cytometer (Beckman-Coulter). Results were analyzed using Cytoexpert 2.5 (Beckman-Coulter) and specific scripts developed using the R programming language.

### Statistics

Statistical analyses were performed using GraphPad Prism software. Data were assessed for normality and homogeneity of variances prior to statistical testing. Comparisons of size between population groups were performed using the Kruskal–Wallis test, followed by Dunn’s post hoc test for multiple comparisons, or the two-tailed Mann–Whitney U test for pairwise comparisons. For forward scatter (FSC) and side scatter (SSC) parameters, Welch’s ANOVA followed by Dunnett’s T3 post hoc test was used for multiple-group comparisons, while the two-tailed unpaired Student’s *t*-test was used for pairwise comparisons. Statistical significance was set at *P* < 0.05, with significance levels indicated as follows: **** *P* < 0.0001, *** *P* < 0.001, ** *P* < 0.01, and * *P* < 0.05.

## ACKNOWLEDGMENTS

This work was supported by the Agence Nationale de la Recherche (ANR) under grant ANR-22-CE44-0005 (PTMyco project) and from a specific support of I2BC through the Myosotis internal program. We also acknowledge the sequencing and bioinformatics expertise of the I2BC High-Throughput Sequencing Facility, supported by France Génomique (funded by the French National Program “Investissements d’Avenir”, ANR-10-INBS-09). This work also benefited from the Imagerie-Gif core facility, supported by the Agence Nationale de la Recherche (FBI ANR-24-INBS-0005 [BIOGEN]; SPS ANR-17-EUR-0007, EUR SPS-GSR), with additional financial support from ITMO Cancer of Aviesan and INCa, on funds administered by Inserm. We acknowledge financial support from the CNRS-CEA “METSA” French network (FR CNRS 3507) on the plateform LPS_CRYO. We further acknowledge the facilities and expertise of MIMA2 (Microscopy and Imaging Facility for Microbes, Animals and Foods; Université Paris-Saclay, INRAE, AgroParisTech, 78350 Jouy-en-Josas, France; DOI: 10.15454/1.5572348210007727E12).

We are particularly grateful to C. Regeard (I2BC) and Jéril Degrouard (LPS) for introducing us to the CEMOVIS facilities at LPS. Finally, we would like to thank all the members of the BacSurf team in the Department of Microbiology at I2BC and more particularly Christiane Dietrich, for their valuable support and insightful discussions throughout this project.

