## Supplementary material for "Uncoupling mycomembrane biogenesis from mycolic acid synthesis reveals a distinct role for mycoloyltransferases in mycobacterial cell division": Fig S1-S8

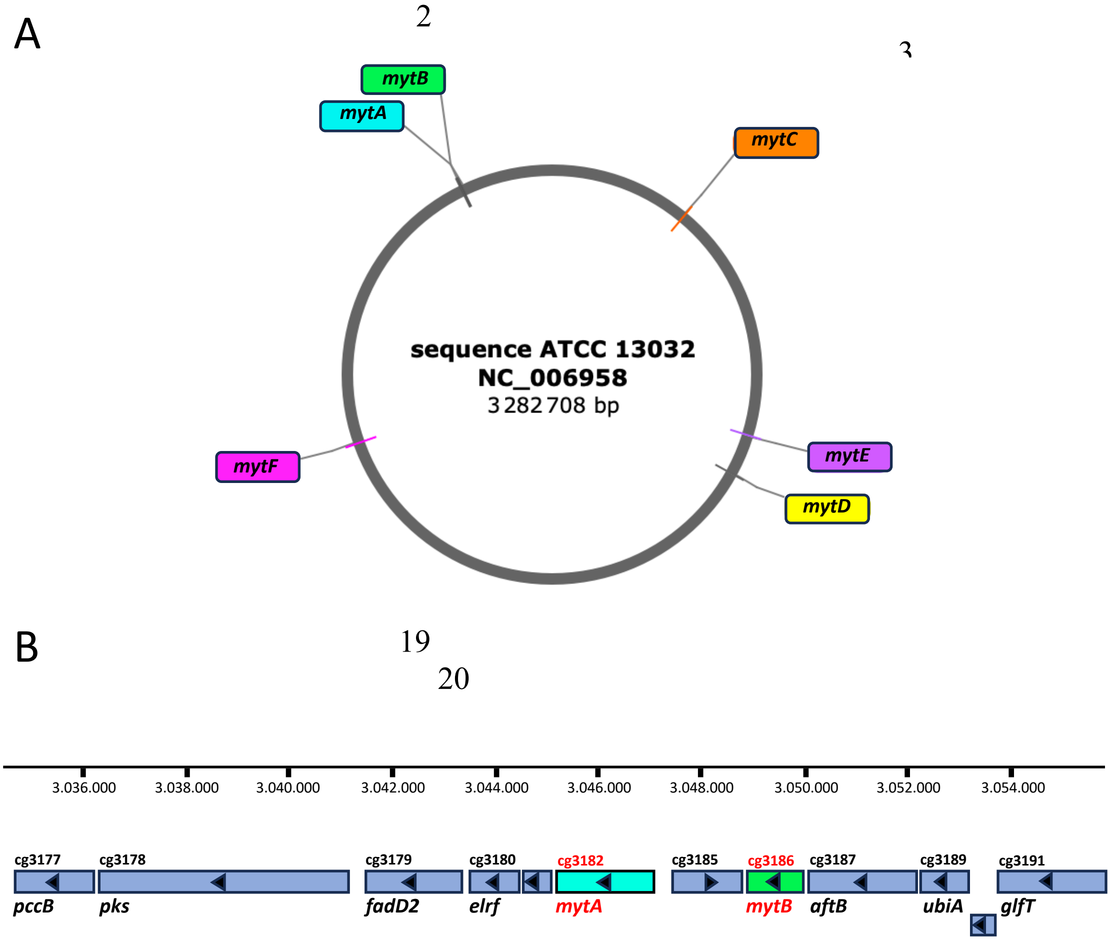


**Figure S1: Genes encoding mycoloyltransferases in *C. glutamicum*. (A)** Distribution of *myts* genes in *C. glutamicum* ATCC13032 genome. *mytC*: *cg0413*; *mytE*: cg1052; *mytD*: *cg1170*; *mytF*: *cg2394*; *mytA*: *cg3182*; *mytB*: *cg3186.* **(B)** Region containing *mytA* and *mytB* genes, and highly conserved among Mycobacteriales. Cg3177/PccB: propionyl-CoA carboxylase beta chain; Cg3178/Pks: polyketide synthase; Cg3179/FadD2: fatty acyl-AMP ligase; Cg3180/Elrf: envelope lipids regulation factor; Cg3181: hypothetical secreted protein; Cg3182/MytA: mycoloyltransferase; Cg3185: hypothetical protein; Cg3186/MytB: mycoloyltransferase; Cg3187/AftB: arabinosyltransferase; Cg3189/UbiA: prenyltransferase; Cg3190: 5'-phosphoribosyl-monophospho-decaprenol phosphatase; Cg3191/Glft: galactosyltransferase.

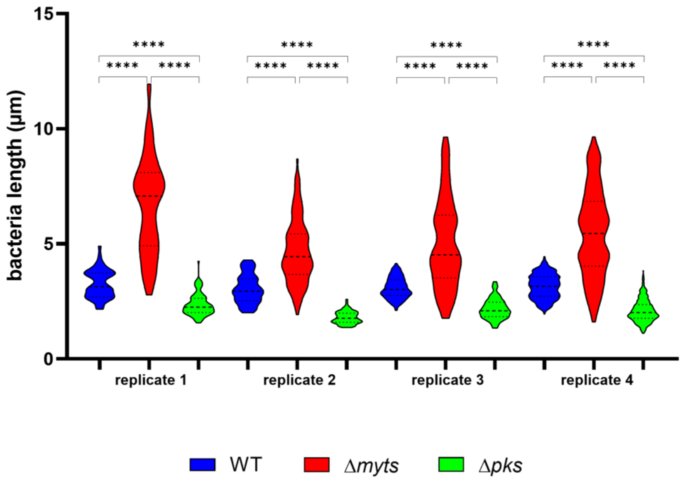


**Figure S2: Violin plots of WT, Δ*myts* and Δ*pks* cell lengths. Violin plots of WT, Δ*myts* and Δ*pks* cell lengths.** Violin plots illustrating the distribution of cell lengths measured for each strain in each biological replicate during the exponential growth phase at 25 °C. Cell length differed significantly between populations within each biological replicate (Kruskal-Wallis test,****P<0,0001). All pairwise comparisons were highly significant (Dunn's post-hoc test,****, P < 0.0001). **(see Table S3)**.

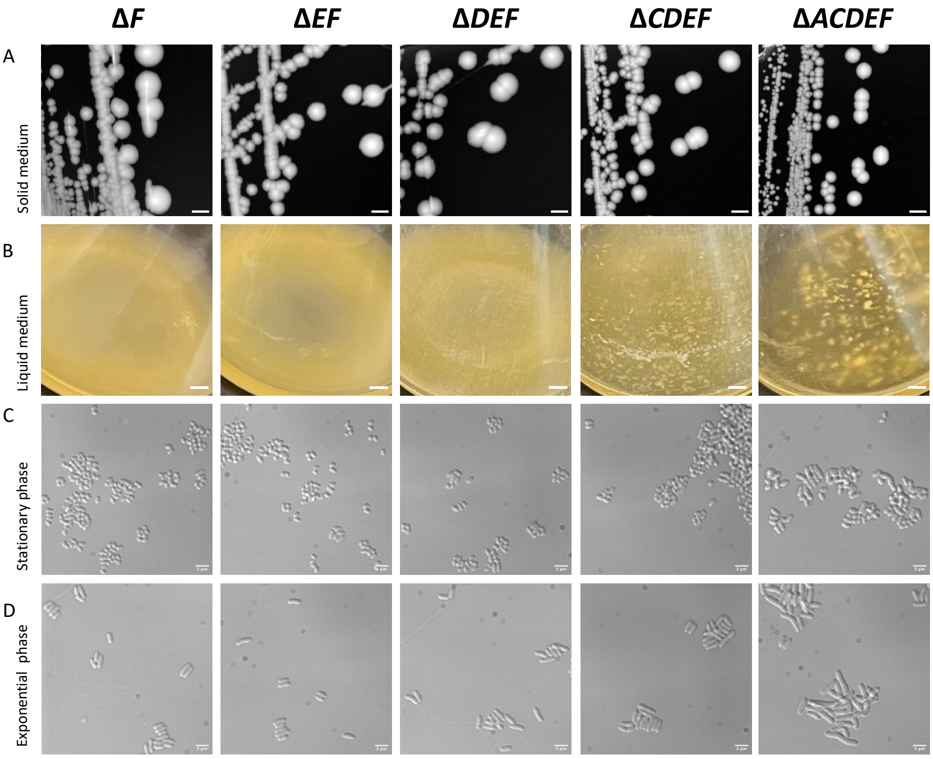


**Figure S3: Morphology and agregation of intermediate mutants. (A)** Colony morphology on solid BHI medium (arbitrary scale). **(B)** Appearance of cultures grown in liquid BHI medium for 24 h at 30°C (arbitrary scale). **(C-D)** Representative differential interference contrast (DIC) micrographs of cells grown in BHI medium during stationary phase and exponential phase (scale bars, 3 µm).

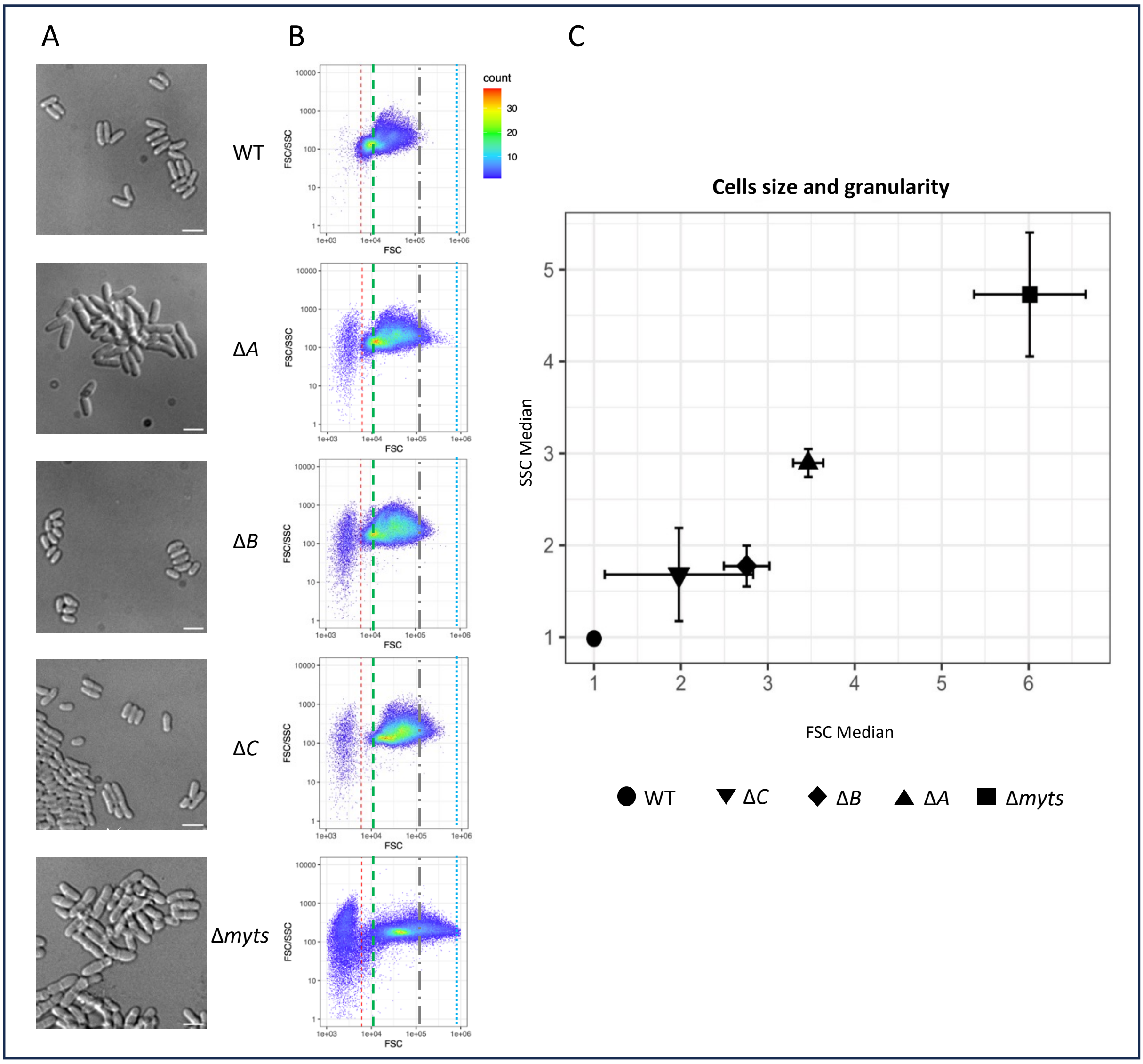


**Figure S4: Morphological characteristics of Δ*myts and single* Δ*A,* Δ*B,* Δ*C strains*. (A)** Representative differential interference contrast (DIC) micrographs of cells grown in BHI medium at exponential phase (scale bars, 3 µm). **(B)** Characteristic population profiles illustrated by scatter plots showing the SSC/FSC ratio as a function of FSC for the replicates closest to both the median and the mean. The dotted red line indicates the FSC threshold above which values were included in the median calculations. The dashed green line marks the maximum FSC value for WT and the begining of the corresponding range for the mutant. The grey dashed-and-dotted line indicates the maximum FSC value for the mutant, while the dotteded-blue line marks the upper limit of this range. **(C)** Scatter plot of flow cytometry data showing the median SSC value as a function of the median FSC value for two independent biological replicates normalized to WT.

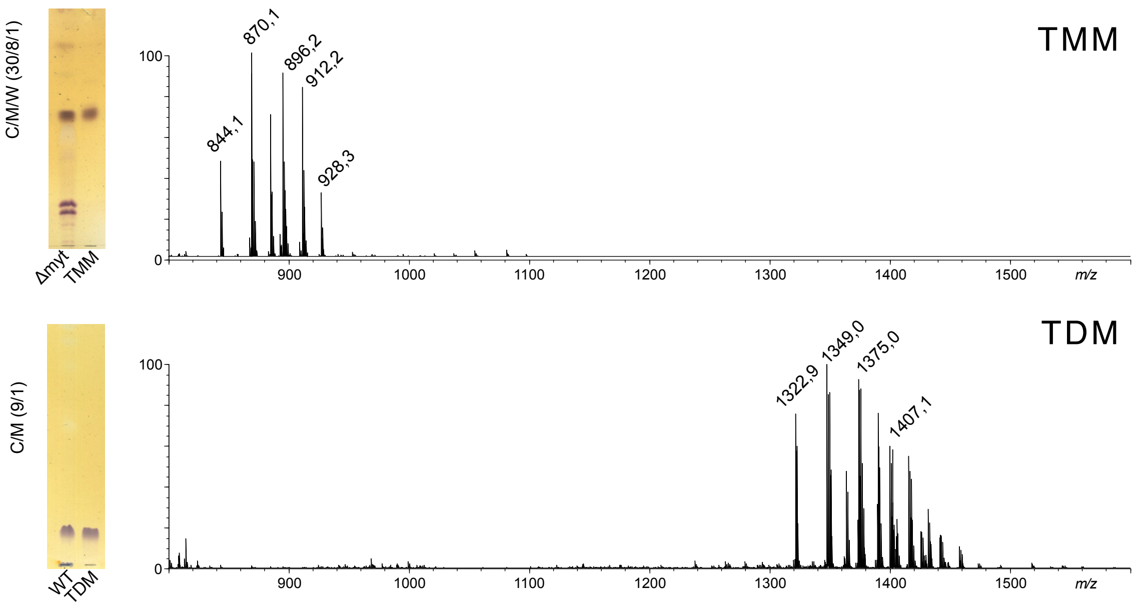


**Figure S5: Analysis of TMM and TDM**.
The two bands at retention factor (Rf) at 0.46 and 0.92, tentatively identified as TMM and TDM respectively based on previous reports, were purified from WT and Δ*myts* strains by silica gel chromatography and individually analyzed by mass spectrometry (MS). TLC analysis confirmed the purity of both compounds (left panels). Their MALDI-MS analysis showed complex patterns of multiple signals ranging from m/z 844 to 928 and from *m/z* 1322 and 1459 assigned to di-hexosides (Hex2) substituted by one or two mycolic acids, respectively, in agreement with their identification as TMM and TDM (Table S4). Each signal pattern corresponds to distinct molecular species of TDM and TMM, comprising mycolic acids that vary in carbon chain length and degree of unsaturation. Our data align well with previous reports, confirming that the predominant corynomycolic acids in TDM are C32:0, C34:1, and C36:2, and that mycolic acids with odd carbon numbers are present in *C. glutamicum* [1]. C/M, chloroform /methanol and C/M/W, chloroform/methanol/water.

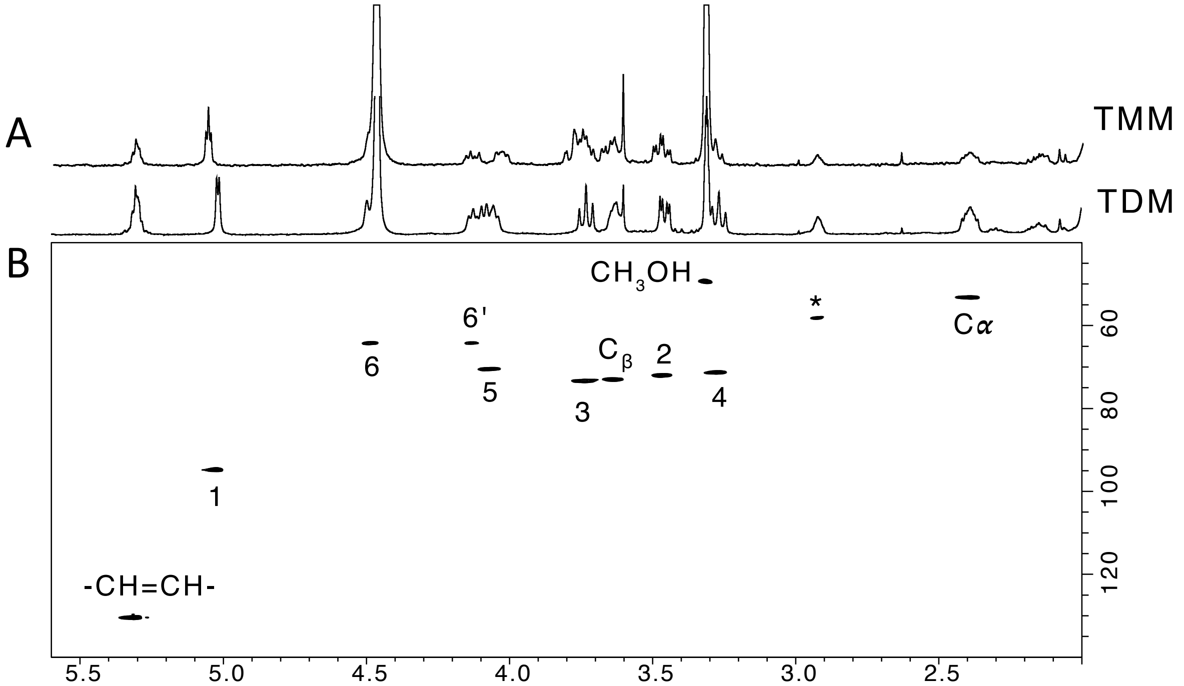


**Figure S6: Analysis of TMM and TDM by NMR**. The structure of purified TMM and TDM was confirmed by **(A)** $^{1}$H 1D and **(B)** $^{1}$H/$^{13}$C-HSQC NMR spectroscopy. TMM and TDM showed very similar $^{1}$H-NMR spectra in which all signals from the trehalose could be assigned. Strong deshielding of H6,6' signals clearly demonstrates that both C6 of the trehalose moiety are mycoloylated in TDM. It should be noted that signals of both glucose residues of the TDM. including the anomers, are perfectly superimposed due to the molecular symmetry. In contrast. the $^{1}$H spectrum of TMM is not, as demonstrated by the dispersity of the anomeric signals that forms a pseudo triplet instead of a perfect doublet as for TDM. 1-6 refer to the H/C positions of glucose residues, Ca and Cb to the carbons next to the CO function of mycolic acids and CH=CH to the signals of insaturations. *unidentified signal.

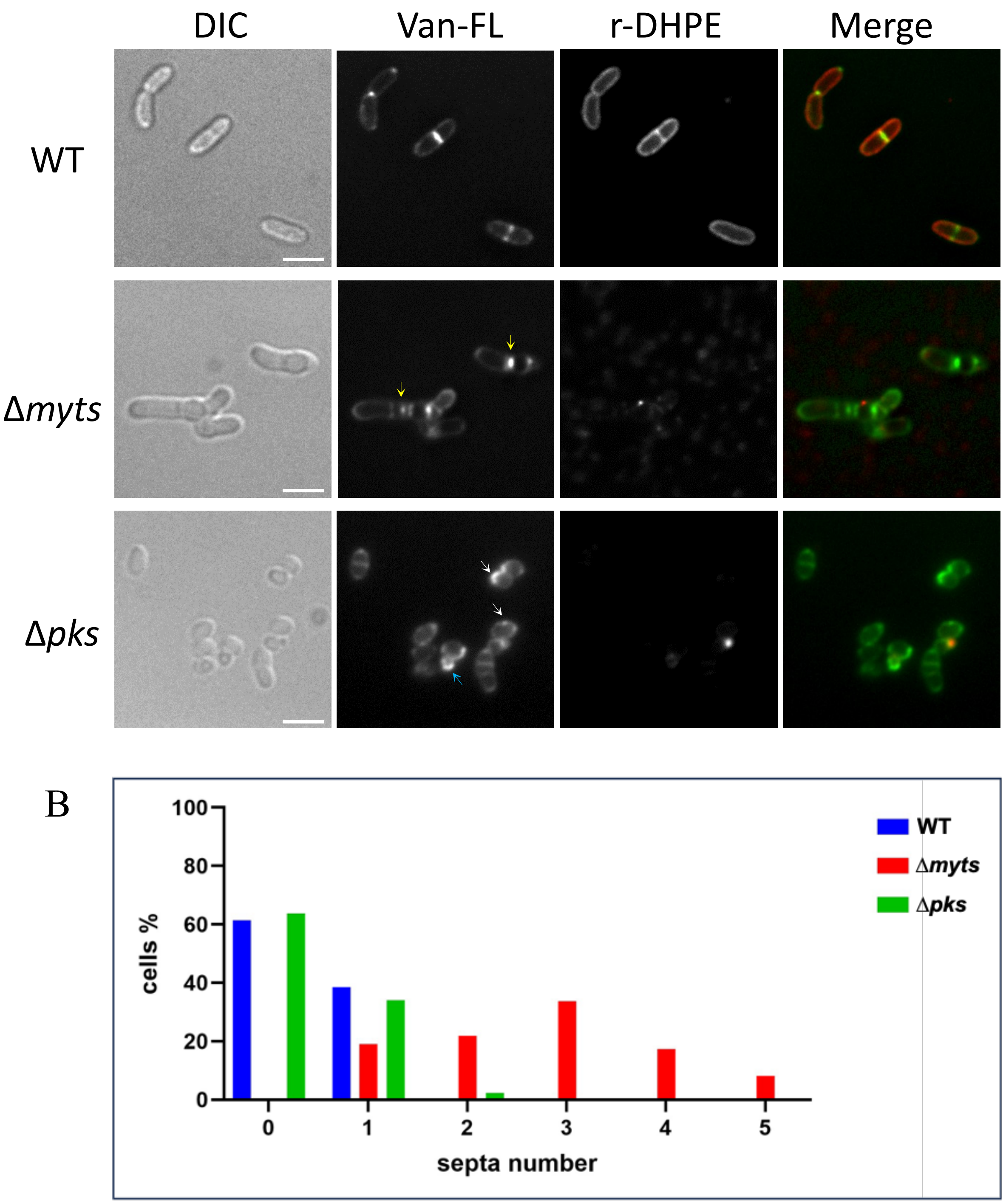


**Figure S7: Fluorescent labeling of peptidoglycan and mycomembrane in the different strains. (A)** Differential interference contrast (DIC) and fluorescence micrographs of bacterial cells, treated with Van-FL or rDHPE during the exponential phase. The left panel shows a merge of the two channels (peptidoglycan in green and cell membrane in red). Yellow arrow: areas of increased intensity along the Van-FL stained septum. White arrow: polar and non-polar areas showing more intense staining with Van-FL. Blue arrow: bud-like structures. The microscopy images were acquired using identical exposure times and dynamic range settings, ensuring direct comparability between conditions. **(B)** Vancomycin staining was used to assess the number of septa in approximately 250 cells per population and determine the percentage of cells with different numbers of septa.

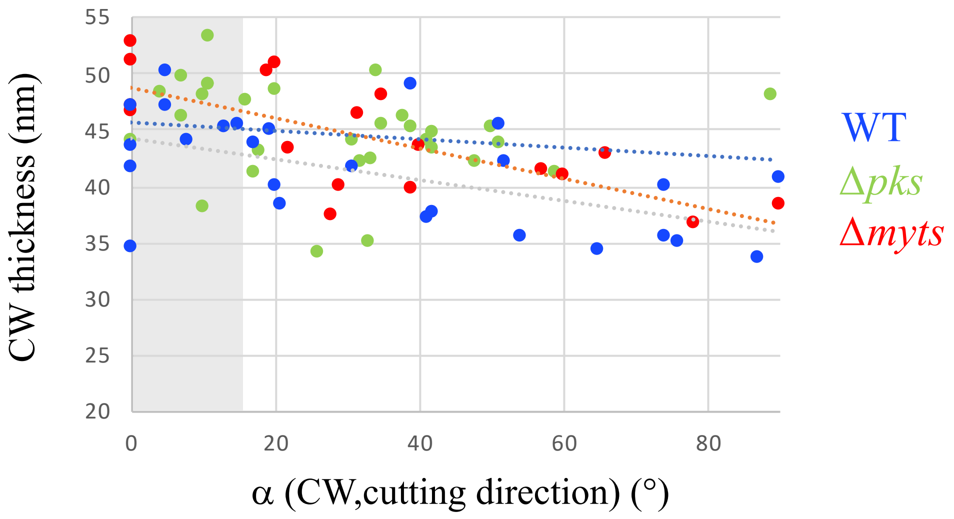


**Figure S8: Measurement of cell wall thickness**. Cryo-sectioning is associated to compression, maximal in the cutting direction [2], [3]: the measured CW thickness decreases with the increase of its angle to the cutting direction. Reliable measurements (reported in Figure 3G) are therefore obtained where the CW is parallel or close to the cutting direction ($\alpha =$ 0-15 °, shaded in grey above): 44.2 ± 4.4 nm (WT), 47.2± 4.3 nm (Δ*myts*), 49.3 ± 2.9 nm (Δ*pks*).

[1] P. Rath *et al.*, “Cord factor (trehalose 6,6′-dimycolate) forms fully stable and non-permeable lipid bilayers required for a functional outer membrane,” *Biochim. Biophys. Acta Biomembr.*, vol. 1828, no. 9, pp. 2173–2181, 2013, doi: 10.1016/j.bbamem.2013.04.021.

[2] A. ALAMOUDI, D. STUDER, and J. DUBOCHET, “Cutting artefacts and cutting process in vitreous sections for cryo-electron microscopy,” *J. Struct. Biol.*, vol. 150, no. 1, pp. 109–121, Apr. 2005, doi: 10.1016/j.jsb.2005.01.003.

[3] A. Leforestier, N. Lemercier, and F. Livolant, “Contribution of cryoelectron microscopy of vitreous sections to the understanding of biological membrane structure,” *Proceedings of the National Academy of Sciences*, vol. 109, no. 23, pp. 8959–8964, Jun. 2012, doi: 10.1073/pnas.1200881109.
