## Supplementary material for "Uncoupling mycomembrane biogenesis from mycolic acid synthesis reveals a distinct role for mycoloyltransferases in mycobacterial cell division": Fig. S9-S17

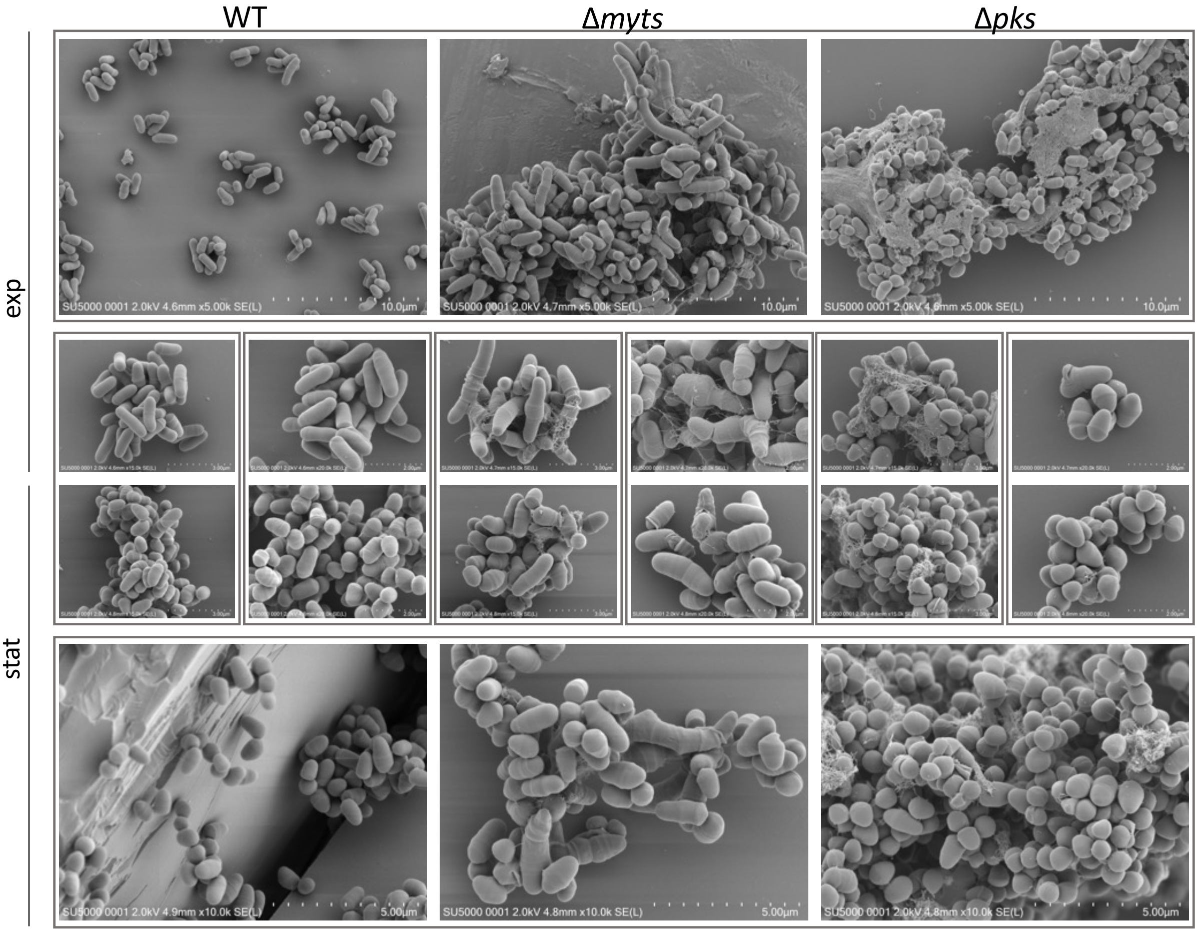


**Figure S9: Observation of cell morphology by scanning electron microscope (SEM).** For direct comparison between strains. cells are presented side by side in exponential **(top)** and stationary **(bottom)** phases, with consistent magnification within each phase (scale bars: 10 µm and 5 µm. respectively). To compare growth phases, images acquired at the same magnification (scale bars: 3 µm and 2 µm) are shown for each strain **(central panels).**

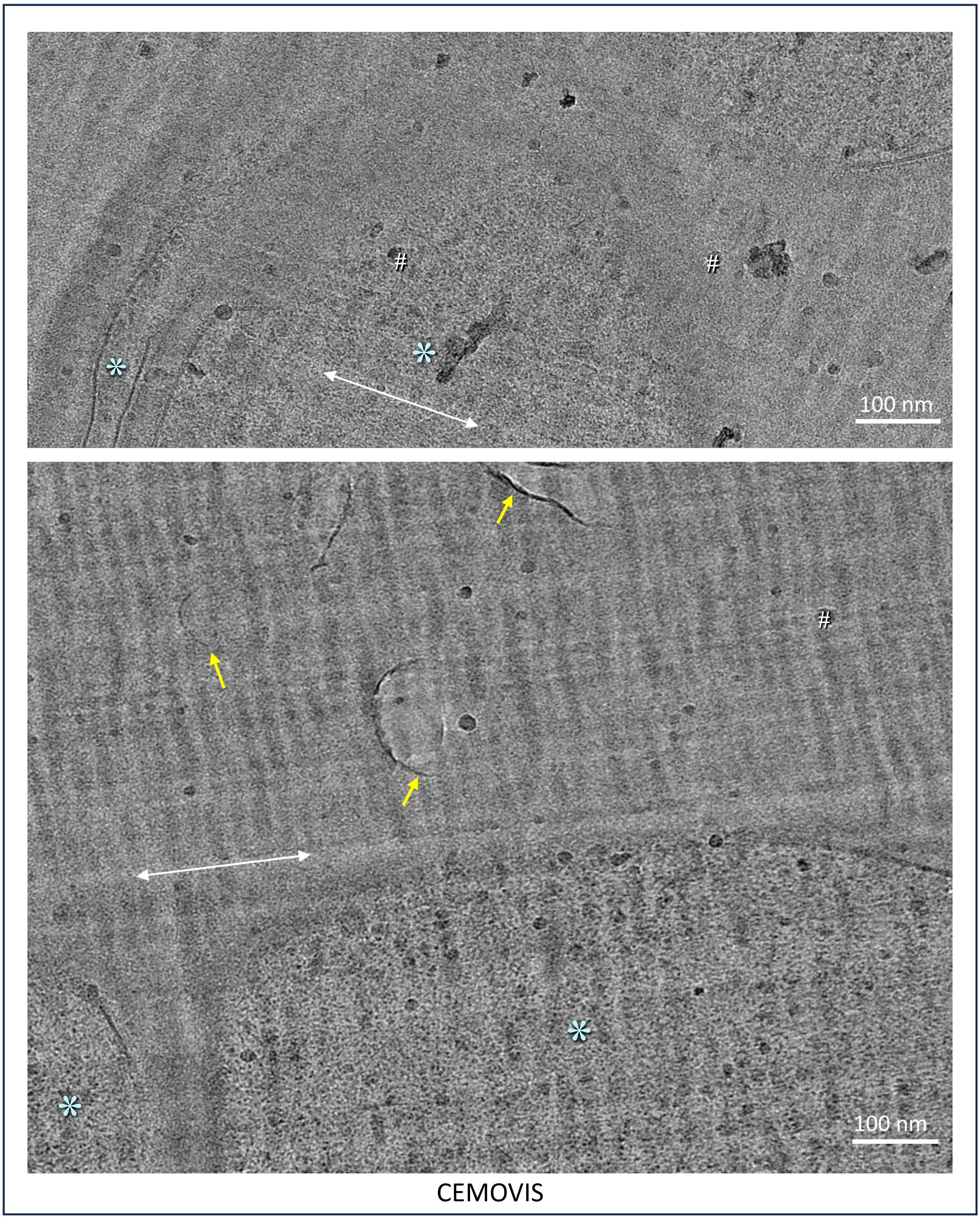


**Figure S10: CEMOVIS observations of Δ*myts* cells** (complementary data for Figure 4). Overall views of the cell reveal series of septa separating compartments (*). Membrane debris are observed within the surrounding medium (yellow arrows). The white double arrow underlines knife marks on the section's surface, highlighting the cutting direction. # indicates small ice contaminations deposited on the section's surface.

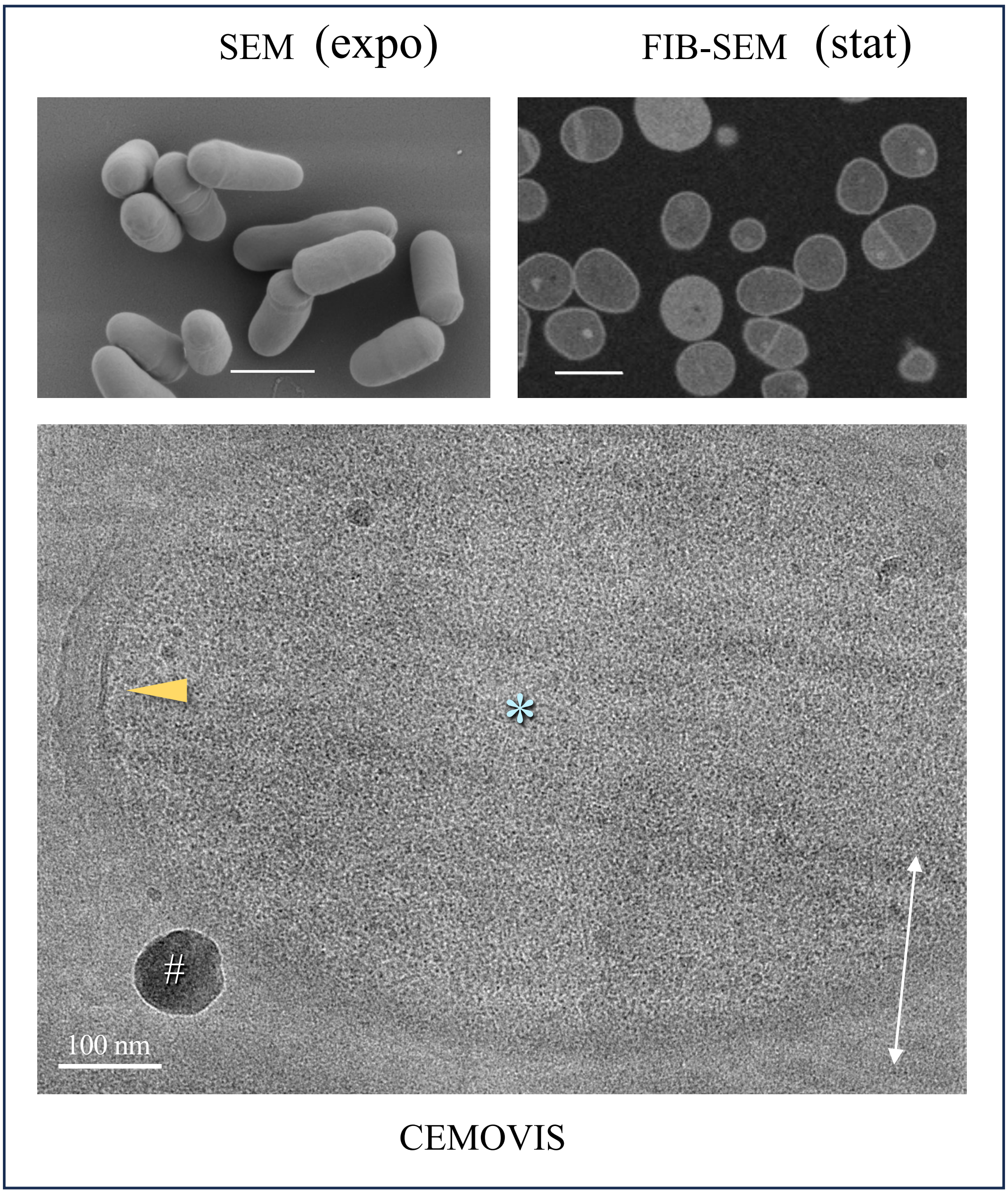


**Figure S11: Electron microscope observations of WT cells** (complementary data for Figure 4). SEM in the top left panel, the characteristic homogeneous rod-shaped morphology of *C. glutamicum* in the exponential phase. FIB-SEM in the top right panel, the characteristic coccoid morphology of WT in the stationary phase is observed. Note the very clean background of these images, in contrast to that of Δ*pks* or Δ*myts* in Figure 4. In the bottom panel, CEMOVIS images of bacteria derived from colonies exhibiting an homogeneous cytoplasm (asterisk) and regular cell wall with its two membranes (arrowhead). The white double arrow indicates the cutting direction. # indicates an ice crystal contamination deposited on the section's surface. SEM and FIB-SEM scale bars: 1 µm.

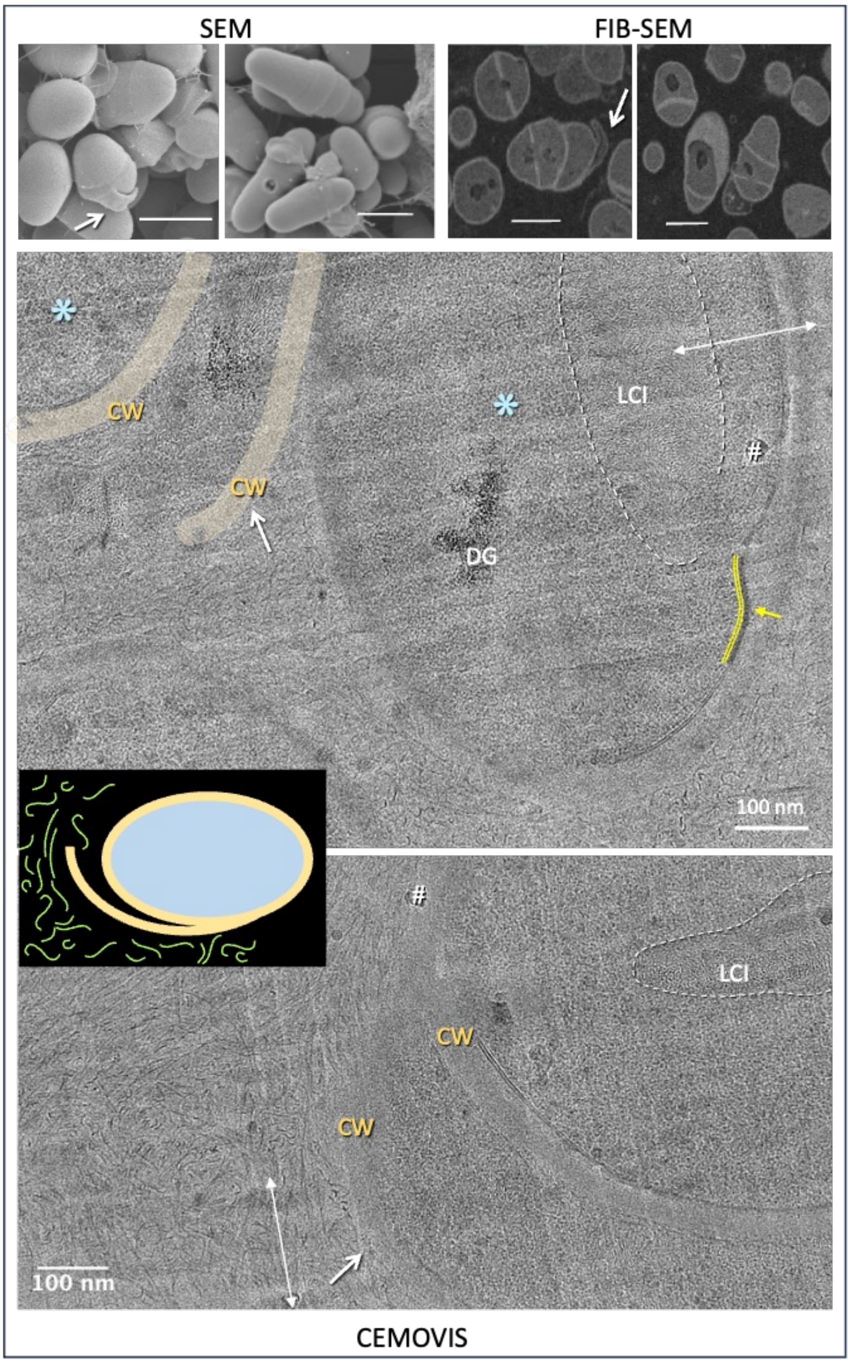


**Figure S12: Electron microscope observations of Δ*pks* cells** (complementary data for Figure 4). In the top left panels, SEM images representative of the two morphological types of Δ*pks* (round or slightly elongated and stocky) observed during the exponential phase. The corresponding FIB-SEM images from the stationary phase are shown in the top right panels. The 'hood'-shaped structures are indicated by white arrows; they are also visible in the CEMOVIS images (cells from colonies) in the central panel. Deformations of the cell surface are underlined (yellow arrows). The white double arrow indicates the cutting direction. # indicates an ice crystal contamination deposited on the section's surface and * the cytoplasm. DG: dense granules. LCI: liquid crystalline inclusion typical of nucleoïd [1], [2]. Insert: sketch of a Δ*pks* cell, with an open cell wall cap surrounded by numerous filaments.

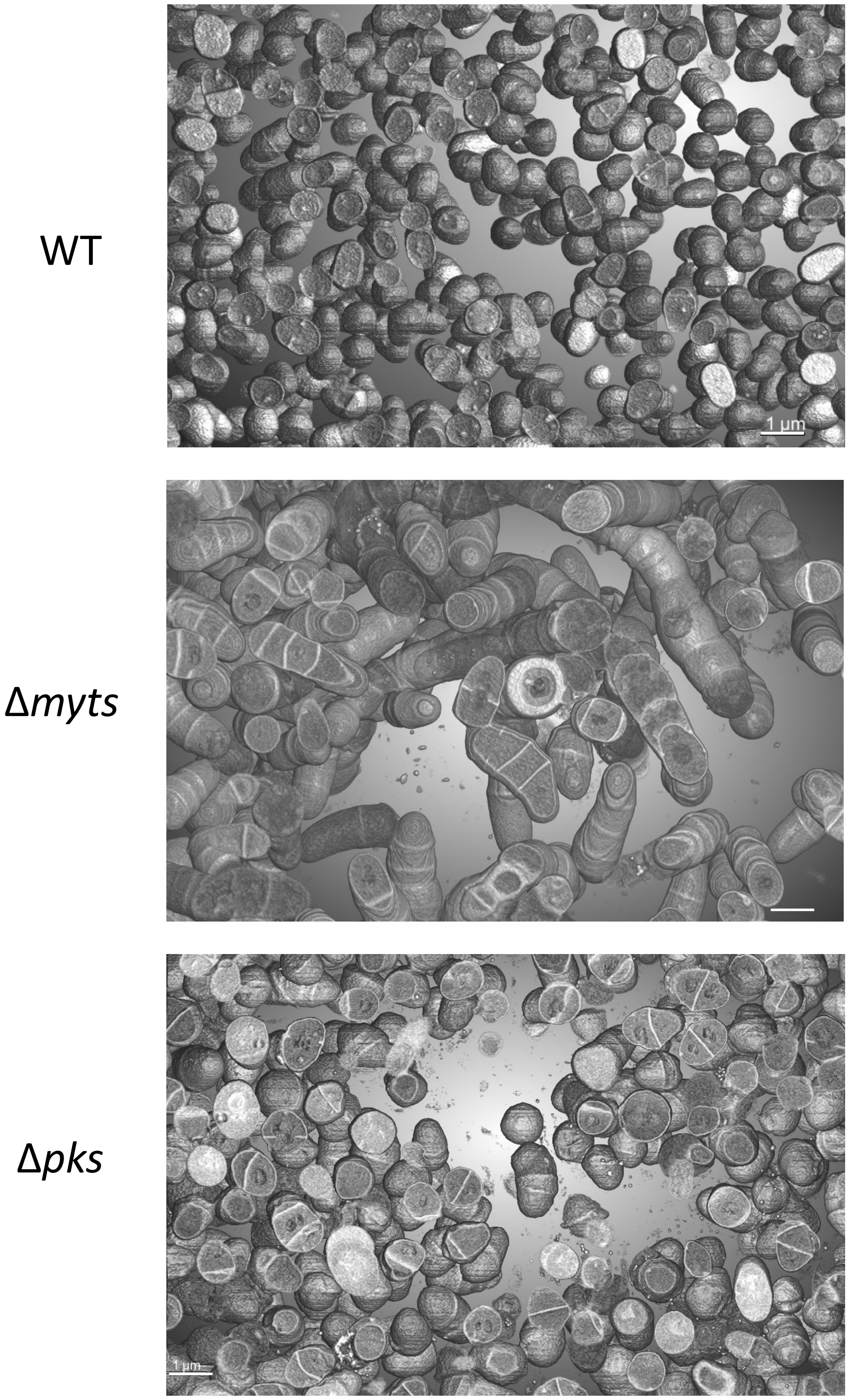


**Figure S13: 3D FIB-SEM images of strains.** Snapshot of 3D volumetric data obtained via FIB-SEM for WT, Δ*myts* and Δ*pks* in stationary phase, with a resolution of 25 nm/pixel. The cells display the morphological features characteristic of this phase: coccoid for WT with rare individual septa, elongated with multiple septa for Δ*myts*, and rounded and stocky with one or two septa for Δ*pks*.

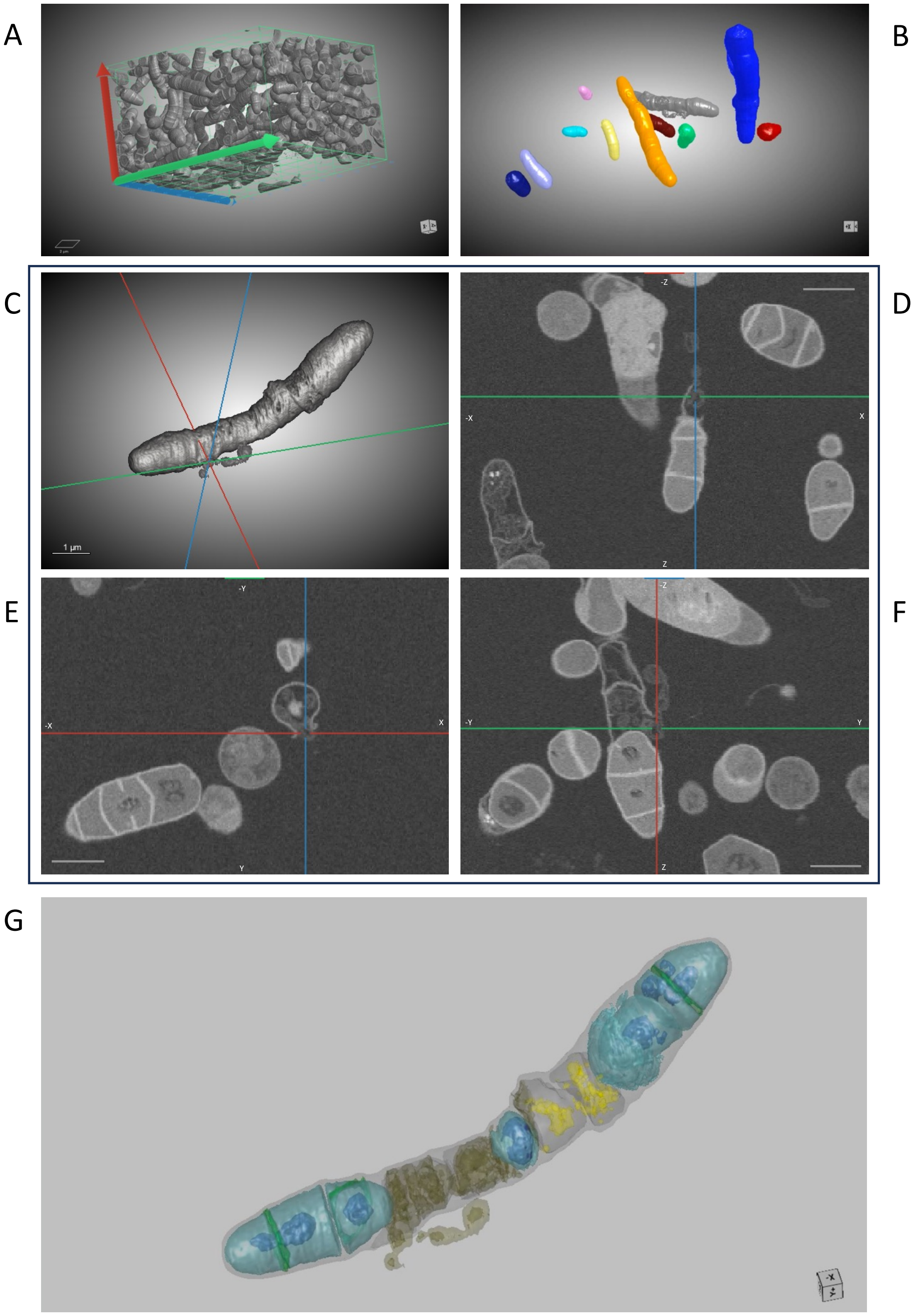

A
B
C
D
E
F
G
1 µm
-Z
-X
X
Z
-Y
Y

**Figure S14: FIB-SEM imaging and 3D reconstruction of the Δmyts mutant.**

**(A)** Three-dimensional rendering of the resin-embedded strain Δ*myts* from the FIB-SEM image stack. The image is oriented with the x-axis (red), y-axis (blue), and z-axis (green). Scale bar, 2 µm. **(B)** Three-dimensional rendering of 11 randomly selected bacteria based on the segmentation of FIB-SEM images. The maximum Feret diameter of each bacterium is indicated: gray (7.84 µm), brown (43.56 µm), green (3.66 µm), orange (8.67 µm), yellow (3.95 µm), purple (3.42 µm), pink (2.34 µm), light blue (2.15 µm), navy blue (2.02 µm), royal blue (5.60 µm), and red (1.63 µm). See Video 3. **(C)** Three-dimensional rendering of a representative segmented bacterium of the Δ*myts* mutant (in gray), shown with corresponding 2D FIB-SEM cross-sections (**(D)** XZ, **(E)** XY, and **(F)** YZ) aligned along the spatial axes **(G)** Three-dimensional model of a representative bacterium of the Δ*myts* mutant (the gray), reconstructed from the segmentation of the corresponding raw 2D FIB-SEM cross-section images. Cellular structures are segmented as follows: cell wall (gray), cytoplasm (cyan), DNA (navy blue), debris (yellow), and vesicles (khaki). Scale bars, 1 µm. The reconstructed bacterium consists of 10 compartments separated by 9 complete septa. Four compartments are lysed, with their envelopes containing holes or large openings oriented perpendicular to the septa. These compartments have lost their cytoplasm, leaving only cellular debris or vesicles that appear to be forming and escaping through the openings. One additional compartment still contains cytoplasm but is completely compressed by the adjacent cell growing within it, resulting in a markedly curved septum. The five remaining viable compartments, enclosed within the same cell envelope, are located at both poles and in the central region. The central compartment is extremely small, whereas the two polar compartments are undergoing division, with incomplete septa forming above the segregating DNA. One of these nascent septa is misaligned. See Video 4.

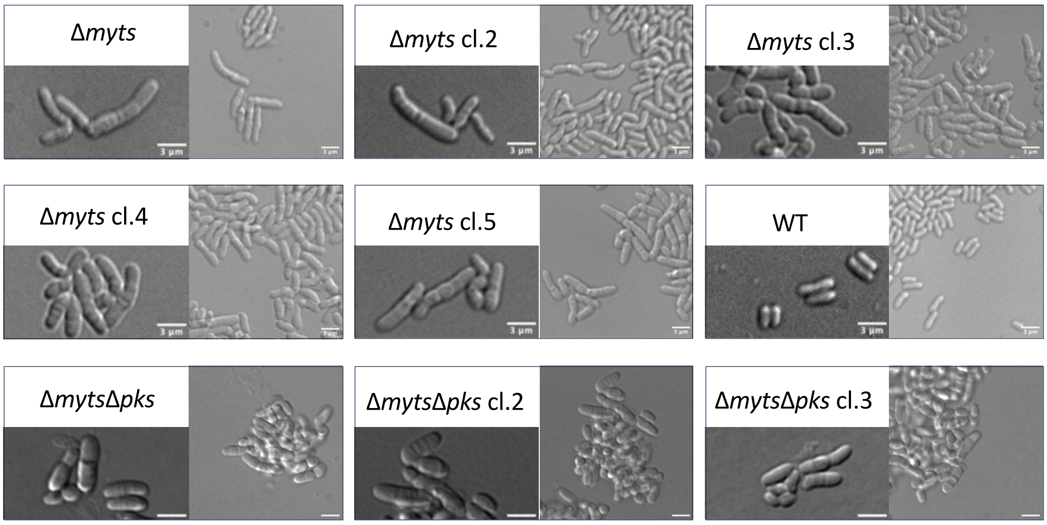


**Figure S15: Morphology of Δ*myts* and Δ*myts*Δ*pks* variants compared to WT.** Representative differential interference contrast (DIC) micrographs of cells at exponential phase in BHI medium (scale bars, 3 µm).

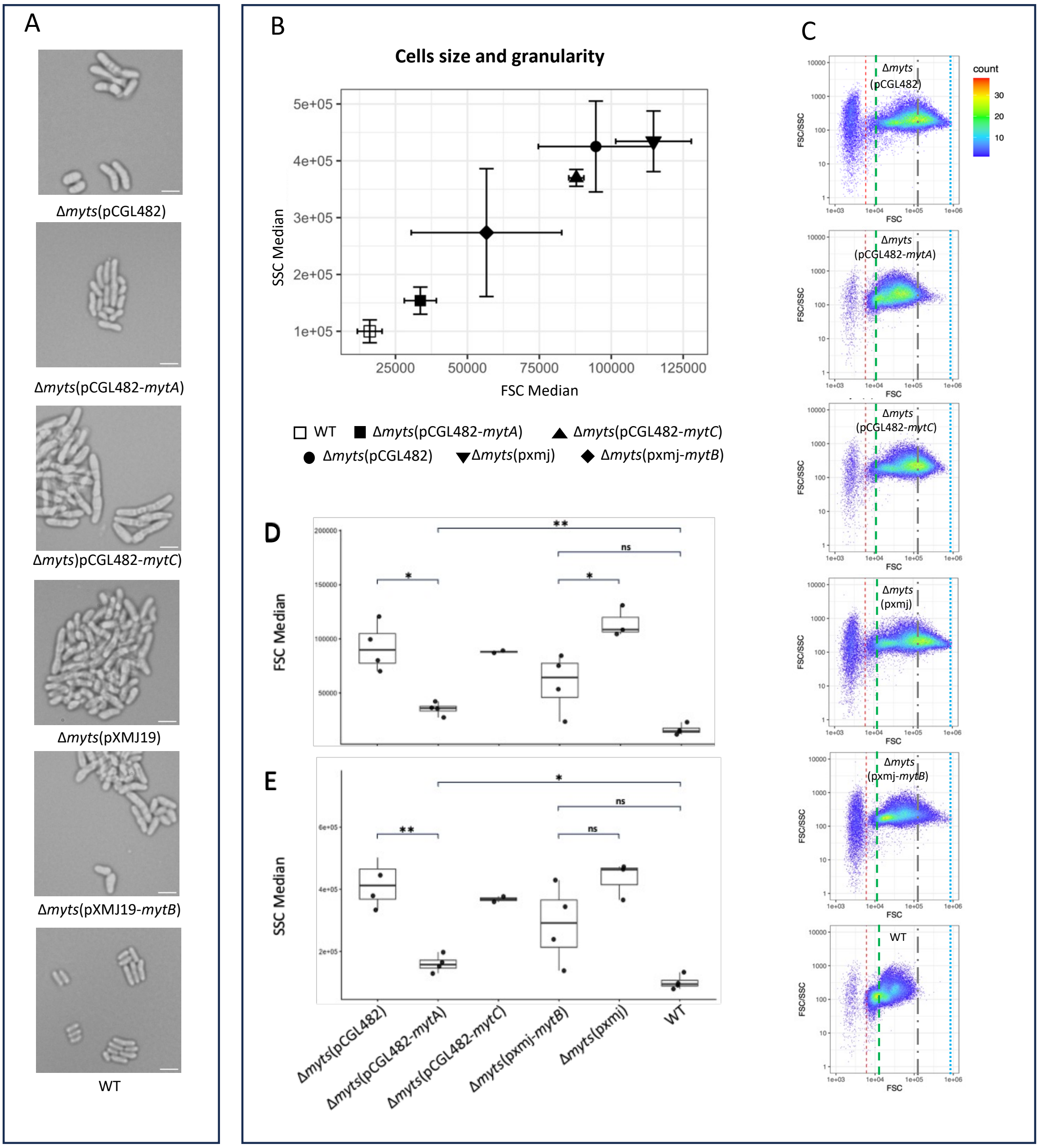


**Figure S16: Morphological characteristics of Δ*myts* complemented strains.** (A) Representative differential interference contrast (DIC) micrographs of cells grown in BHI medium during exponential phase (scale bars, 3 µm). Images were acquired using a CyScope® HP (CY-S-4041_R, Sysmex Partec). (B) Scatter plot showing median SSC as a function of median FSC for individual biological replicates in (C–D). Box plots showing median FSC (C) and SSC (D) values measured by flow cytometry. Values represent the mean of the median values obtained from technical replicates for each biological replicate. Statistical results for *mytC* complementation are not displayed because of the limited sample size (n = 2) and should be interpreted with caution. (E) Representative SSC/FSC profiles from replicates closest to the median and mean. The dotted red line indicates the FSC threshold used for median calculations. The dashed green line indicates the maximum WT FSC value, the grey dashed-dotted line the maximum mutant FSC value, and the dashed blue line the upper limit of the corresponding FSC range.

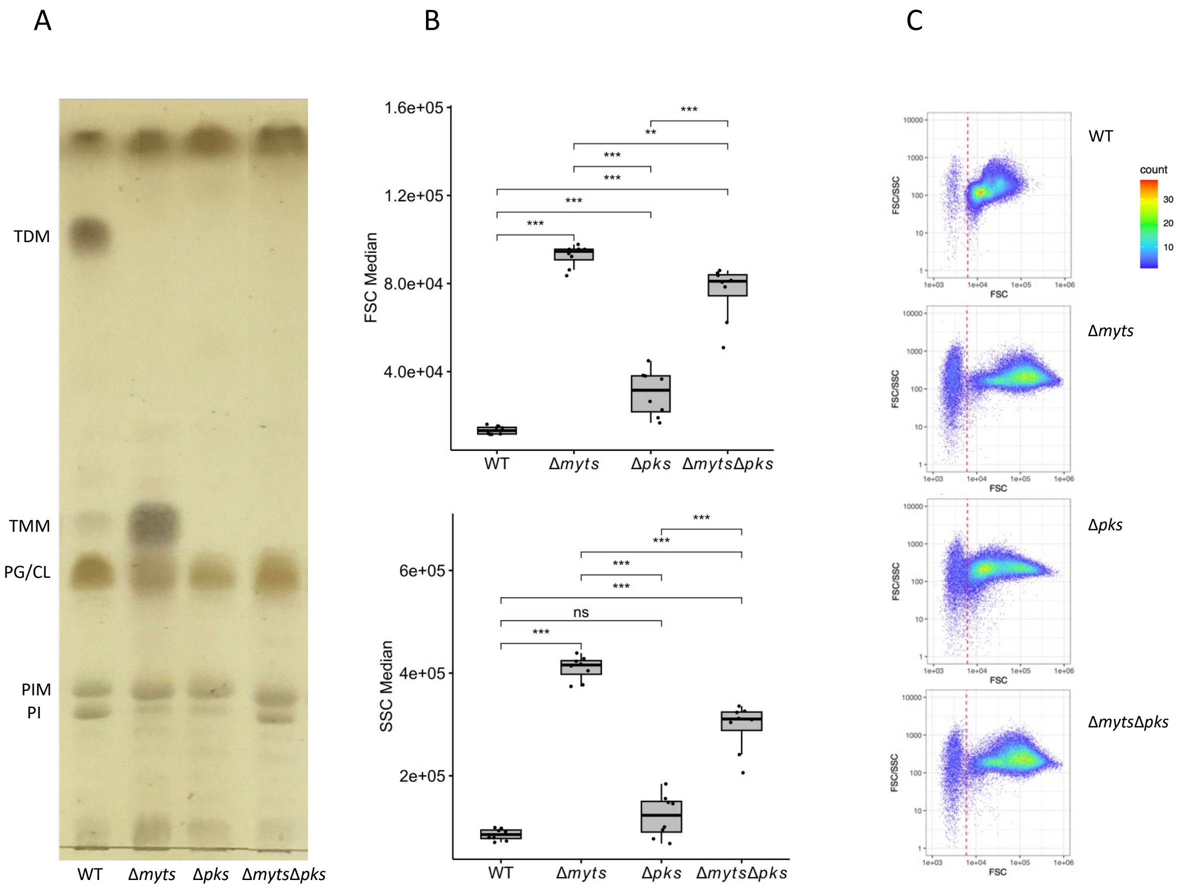


**Figure S17: Lipid composition and morphological characteristics of Δ*myts*Δ*pks* cells. (A)** Thin-layer chromatography analysis of whole cell lipid extracts. **(B)** Box plots showing the median FSC or SSC values obtained by flow cytometry for two technical replicates of four independent biological replicates. **(C)** Characteristic population profiles, illustrated by representative replicate scatter plots showing the SSC/FSC ratio versus FSC. Dotted red line represented threshold on FSC-parameter beyond which the values were considered for median calculations.

[1] M. Eltsov and J. Dubochet, “Fine structure of the Deinococcus radiodurans nucleoid revealed by cryoelectron microscopy of vitreous sections,” *J. Bacteriol.*, vol. 187, no. 23, pp. 8047–8054, Dec. 2005, doi: 10.1128/JB.187.23.8047-8054.2005.

[2] B. Yee, E. Sagulenko, G. P. Morgan, R. I. Webb, and J. A. Fuerst, “Electron tomography of the nucleoid of Gemmata obscuriglobus reveals complex liquid crystalline cholesteric structure,” *Front. Microbiol.*, vol. 3, 2012, doi: 10.3389/fmicb.2012.00326.
