## Supplementary material for "Uncoupling mycomembrane biogenesis from mycolic acid synthesis reveals a distinct role for mycoloyltransferases in mycobacterial cell division": Tables S1-S5

**Table S1: Oligonucleotides used in this study.**

| Name | 5'-3' sequence |
|---|---|
| **mytA-del1-XhoI** | tggagcttctcgaggagttcgg |
| **mytA-del2-NcoI** | atcgcccatggagttttccttc |
| **mytA-del3-RcaI** | actgtcatgaagcgcatctacgcaactc |
| **mytA-del4-BamHI** | attcggatccccagattccttc |
| **mytA-ver1** | attgcagcaggcgctgtccc |
| **mytA-ver2** | ttgagcttgtctacgttgc |
| **mytA-ver3** | cgagggatccatgtgggaggtttttcc |
| **mytA-ver4** | actcattgtgagtgttgatcag |
| **mytB-del1-BamHI** | tggttggatcctcaaaagac |
| **mytB-del2-HindIII** | tacgaagcttggacataattctcctc |
| **mytB-del3-HindIII** | tacgaagcttctagacgtcatgaaac |
| **mytB-del4-XhoI** | agtcctcgagtccgaatccggccgtc |
| **mytB-ver1** | atcctgtcatcttacatgtgg |
| **mytB-ver2** | caacgcgacggcaaagac |
| **mytB-ver3** | aatccgccactccgctgg |
| **mytB-ver4** | agcaccggcataaaactcac |
| **mytC-del1-XhoI** | ttatctcgagcgctgaataggtc |
| **mytC-del2-SacII** | aataccgcggtacgtctttccctc |
| **mytC-del3-SacII** | ttaatccgcggatcttggaccac |
| **mytC-del4-BglII** | acggagatctaaaatacggtcg |
| **mytC-ver1** | aattgcgatgtccaccattg |
| **mytC-ver2** | tacccagtcggtgtagtagg |
| **mytC-ver3** | tttgcagtcctcttgcag |
| **mytC-ver4** | ttcctaccaccatgtttg |
| **mytD-del1-BglII** | taatagatctgtcaacgaaagacac |
| **mytD-del2-XbaI** | cttgctctagatatgaagagcttc |
| **mytD-del3-XbaI** | ctagtctagagctctttactagagc |
| **mytD-del4-HindIII** | gccaaagcttgcccaaaccaac |
| **mytD-ver1** | ttgaacagggtggacttg |
| **mytD-ver2** | ttccccaagcaaaggaac |
| **mytD-ver3** | acctgacagatagcgtctg |
| **mytD -ver4** | agaaacaacagcggaatacc |
| **mytE-del1-BglII** | tagcagatctgacccactcaaattcc |
| **mytE-del2-XbaI** | cttgctctagactagcaaccgctacc |
| **mytE-del3-XbaI** | ctagtctagattaagccaatcagcagg |
| **mytE-del4-XhoI** | acgtctcgagatgccattgaac |
| **mytE-ver1** | atgacatcgataagggatg |
| **mytE-ver2** | acaagttcgatgcggcaactc |
| **mytE-ver3** | atcagatcatcgtcgagc |
| **mytE-ver4** | tcataactacttcccttctc |
| **mytF-del1-BglII** | tgctagatcttccactgatcgcagtg |
| **mytF-del2-XbaI** | cttgctctagaaattcctttacgcactg |

| | |
|---|---|
| **mytF-del3-XbaI** | ctcgtctagaggaagcttctacgaatag |
| **mytF-del4-XhoI** | acgtctcgagacgtacggtatataactg |
| **mytF-ver1** | tagcaccggcagxtataatc |
| **mytF-ver2** | tttatgtgattgttgcagc |
| **mytF-ver3** | tgagcatcgcactaccaac |
| **mytF-ver4** | actcttgttccccctaaac |

**Table S2: Statistical analysis of cell lengths for WT, Δ*myts* and Δ*pks* strains.**

| strain | replicate | N cells | mean length | standard deviation | coeff. variation | Min | Max | range | median length |
|---|---|---|---|---|---|---|---|---|---|
| WT | 1 | 46 | 3.259 | 0.6190 | 18.99% | 2.166 | 4.898 | 2.732 | 3.140 |
| | 2 | 71 | 3.042 | 0.6226 | 20.46% | 2.010 | 4.295 | 2.285 | 2.938 |
| | 3 | 227 | 3.078 | 0.4473 | 14.53% | 2.111 | 4.151 | 2.040 | 3.015 |
| | 4 | 762 | 3.153 | 0.5237 | 16.61% | 1.957 | 4.449 | 2.492 | 3.163 |
| | all | 1106 | 3.135 | 0.5216 | 16.64% | 1.957 | 4.898 | 2.941 | 3.117 |
| Δ*myts* | 1 | 63 | 6.729 | 2.121 | 31.52% | 2.782 | 11.93 | 9.151 | 7.077 |
| | 2 | 121 | 4.671 | 1.385 | 29.64% | 1.931 | 8.685 | 6.754 | 4.444 |
| | 3 | 78 | 5.028 | 1.935 | 38.50% | 1.772 | 9.640 | 7.868 | 4.523 |
| | 4 | 278 | 5.467 | 1.890 | 34.57% | 1.611 | 9.645 | 8.034 | 5.451 |
| | all | 540 | 5.372 | 1.914 | 35.62% | 1.611 | 11.93 | 10.32 | 5.104 |
| Δ*pks* | 1 | 97 | 2.357 | 0.4769 | 20.23 | 1.565 | 4.240 | 2.675 | 2.255 |
| | 2 | 94 | 1.798 | 0.2653 | 14.75 | 1.365 | 2.588 | 1.223 | 1.767 |
| | 3 | 83 | 2.176 | 0.4378 | 20.12 | 1.338 | 3.352 | 2.014 | 2.097 |
| | 4 | 411 | 2.096 | 0.4853 | 23.16 | 1.113 | 3.833 | 2.720 | 2.013 |
| | all | 685 | 2.101 | 0.4778 | 22.73% | 1.113 | 4.240 | 3.127 | 2.020 |

**Table S3: Two-tailed Mann–Whitney *p*-values for bacterial length comparisons (related to Figure 1F).**

| **Strains** | | **WT** | | | | **Δ*myts*** | | | | **Δ*pks*** | | | |
|---|---|---|---|---|---|---|---|---|---|---|---|---|---|
| | replicate | 1 | 2 | 3 | 4 | 1 | 2 | 3 | 4 | 1 | 2 | 3 | 4 |
| **WT** | 1 | | ns | ns | ns | | | | | | | | |
| | 2 | | | ns | ns | | | | | | | | |
| | 3 | | | | ns | | | | | | | | |
| | 4 | | | | | | | | | | | | |
| **Δ*myts*** | 1 | **** | **** | **** | **** | | **** | **** | **** | | | | |
| | 2 | **** | **** | **** | **** | | | ns | **** | | | | |
| | 3 | **** | **** | **** | **** | | | | ns | | | | |
| | 4 | **** | **** | **** | **** | | | | | | | | |
| **Δ*pks*** | 1 | **** | **** | **** | **** | **** | **** | **** | **** | | **** | * | **** |
| | 2 | **** | **** | **** | **** | **** | **** | **** | **** | | | **** | **** |
| | 3 | **** | **** | **** | **** | **** | **** | **** | **** | | | | ns |
| | 4 | **** | **** | **** | **** | **** | **** | **** | **** | | | | |

**Table S4: Identification by MALDI-MS of TMM and TDM isolated from Δ*myts* and WT strains. respectively.** *m/z* values were identified in positive mode as $[M+Na]^+$ adducts.

| | *m/z* | Meromycolate | |
|---|---|---|---|
| | M+Na | Carbons | Insaturations |
| TMM | 844.14 | 32 | 0 |
| | 870.18 | 34 | 1 |
| | 886.22 | 35 | 0 |
| | 896.22 | 36 | 2 |
| | 912.26 | 37 | 1 |
| | 928.30 | 38 | 0 |
| TDM | 1323.00 | 64 | 0 |
| | 1349.03 | 66 | 1 |
| | 1375.07 | 68 | 2 |
| | 1401.11 | 70 | 3 |
| | 1407.16 | 70 | 0 |
| | 1427.15 | 72 | 4 |
| | 1433.20 | 72 | 1 |
| | 1459.23 | 74 | 2 |

**Table S5: Synthesis of EM observations (SEM, FIB-SEM, CEMOVIS) of the different strains.**

| Observations | WT | Δ*myts* | Δ*pks* |
|---|---|---|---|
| Agregation | NO | ++ | +++ |
| Multisepta | NO | +++ | + |
| Lysis (empty compartiments) | NO | +++ | +/- |
| External vesicles (mono/multilamellar) | NO | +++ | + |
| External polymers | NO | +/- | +++ |
